# TRAIL agonists rescue mice from radiation-induced lung injury

**DOI:** 10.1101/2023.06.12.544681

**Authors:** Jillian Strandberg, Anna Louie, Seulki Lee, Marina Hahn, Praveen Srinivasan, Andrew George, Arielle De La Cruz, Leiqing Zhang, Liz Hernandez Borrero, Kelsey E. Huntington, Payton De La Cruz, Attila A. Seyhan, Paul P. Koffer, David E. Wazer, Thomas A. DiPetrillo, Christopher G. Azzoli, Sharon I. Rounds, Stephanie L. Graff, Abbas E. Abbas, Lanlan Zhou, Wafik S. El-Deiry

## Abstract

Cancer therapy is often limited by toxicity from pneumonitis. This often-lethal side effect is known to be impacted by innate immunity, and in particular the pathways regulated by the TRAIL death receptor DR5. We investigated whether DR5 agonists could rescue mice from the lethal effects of radiation. We found that two different agonists, parenteral PEGylated trimeric-TRAIL (TLY012) and oral TRAIL-Inducing Compound #10 (TIC10/ONC201), could achieve this goal. Both compounds could completely protect mice from lethality by reducing pneumonitis, alveolar-wall thickness, and oxygen desaturation. At the molecular level, this protection appeared to be due to the inhibition of CCl22, a macrophage-derived chemokine previously associated with radiation pneumonitis and pulmonary fibrosis. The discovery that short-term treatment with TRAIL pathway agonists effectively rescues animals from high doses of radiation exposure has important translational implications.

**One Sentence Summary:** Prevention of lethality, pneumonitis, lung fibrosis and skin dermatitis post-γ-irradiation by short-term treatment with innate immune TRAIL pathway agonists

## INTRODUCTION

It is not sufficient to have effective cancer therapy if there are significant toxicities that impact on quality of life and cause morbidity. A general goal in medicine has been to develop better tolerated therapeutics. Radiation therapy has proven efficacy in local control of cancer as well as patient survival outcomes in multiple tumor types. However, toxicity towards normal tissues has been a challenge in patient treatment. Other cancer therapeutic agents such as bleomycin or immune checkpoint blockade cause morbidity, can interfere with continuing treatment due to lung inflammation and injury, and ultimately contribute to mortality.

In the lungs the so-called late effects of radiation involve inflammation and fibrosis and cause morbidity limiting use of radiation or other cancer therapies in patients who suffer with symptoms (*1, 2*). Radiation-induced pneumonitis develops in ∼10-30% of thoracic cancer patients (*3*). While high-dose radiotherapy to lungs can benefit patients with lung cancer, dose-fractionation is inconvenient and is associated with severe toxicity (*4–6*). Radiotherapy is often combined with immunotherapy including immune checkpoint blockade to treat cancer and this greatly increases the risk and incidence of serious lung injury (*7, 8*). Urgent strategies are needed to reduce the serious toxicities that limit cancer treatment in order to improve the care of patients. There is also a need to develop radiation counter-measures for use in the unfortunate event of a “dirty bomb,” nuclear power plant leak or other unanticipated exposure to high levels of radiation (*9, 10*). Such strategies could be employed to achieve radioprotection if the agents are effective when used post-radiation exposure.

We discovered TRAIL death receptor DR5 as a direct transcriptional target of the p53 tumor suppressor protein (*11, 12*). The p53 protein is central to the cellular DNA damage response (DDR) inflicted by radiation damage. Stabilization of p53 protein after DNA damage activates genes encoding proteins such as p21(WAF1) that mediate cell cycle arrest and repair of DNA damage (*13, 14*). Repair of damage is essential for cell survival without cancer. DR5 is involved in cell death that occurs after chemotherapy or radiotherapy of tumors. The TRAIL pathway that activates DR5 is part of the host innate immune system that suppresses cancer and metastases (*15*).

We previously discovered that *DR5* (also known as *TRAIL-R2 or TNFRSF10b*) gene deletion in mice reduces cell death in multiple organs after lethal γ-irradiation (*16*), and that a sub-lethal dose of whole-body γ-irradiation causes inflammatory lesions and fibrosis in multiple organs including in the lungs of irradiated mice (*17*). This results in lethality of irradiated *DR5*-/- mice 6-8 months post-irradiation. Histologic analysis of lung tissues from *DR5*-/- mice suggests a similarity to what is observed with the late effects of radiation in humans who receive thoracic radiation and develop pneumonitis with deposition of collagen and fibronectin (*17*).

We have known for >15 years that a primary strategy to rescue from the toxic effects of lung radiation might involve treatment with TRAIL pathway agonists which was not particularly feasible or cost-effective years ago due to anticipated duration of treatment of several months. With the availability of novel approaches to stimulation the TRAIL innate immune pathway (*18–21*), we attempted to rescue mice from the toxicities of radiation. Multiple TRAIL innate immune pathway agonists are available including TRAIL-Inducing Compound #10 (TIC10) also known as ONC201 currently in multiple clinical trials for various tumor types (*18, 21, 22*), and TLY012 which is a novel PEGylated trimeric TRAIL formulation that has anti-fibrotic properties (*20*). There are also DR5 agonist antibodies being developed as cancer therapeutics by multiple pharmaceutical companies. Surprisingly, we discovered that short-term treatment with TRAIL pathway agonists for two weeks at or after the radiation exposure effectively rescues mice that receive 20 Gy of thoracic radiation exposure.

## RESULTS

### TRAIL pathway agonists protect mice from lung injury and fibrosis following high-dose thoracic ionizing radiation

We investigated the toxicities of high dose thoracic radiation in wild-type (WT), *DR5*-/- or *TRAIL*-/- C57Bl/6 mice to test predicted outcomes as shown in **Figure 1A**. We hypothesized that *DR5*-/- mice would have more severe radiation-induced pneumonitis and wouldn’t be rescued by either TRAIL ligand or small molecule TRAIL inducing compound TIC10/ONC201 because the receptor for TRAIL (*DR5*) is deleted in the DR5-/- mice (*17*). We also hypothesized that TRAIL formulation TLY012 would rescue pneumonitis in TRAIL-/- mice whereas *TRAIL* gene-inducing compound TIC10/ONC201 would not because the *TRAIL* gene that is upregulated by the drug is deleted in TRAIL-/- mice.

**Figure 1.**
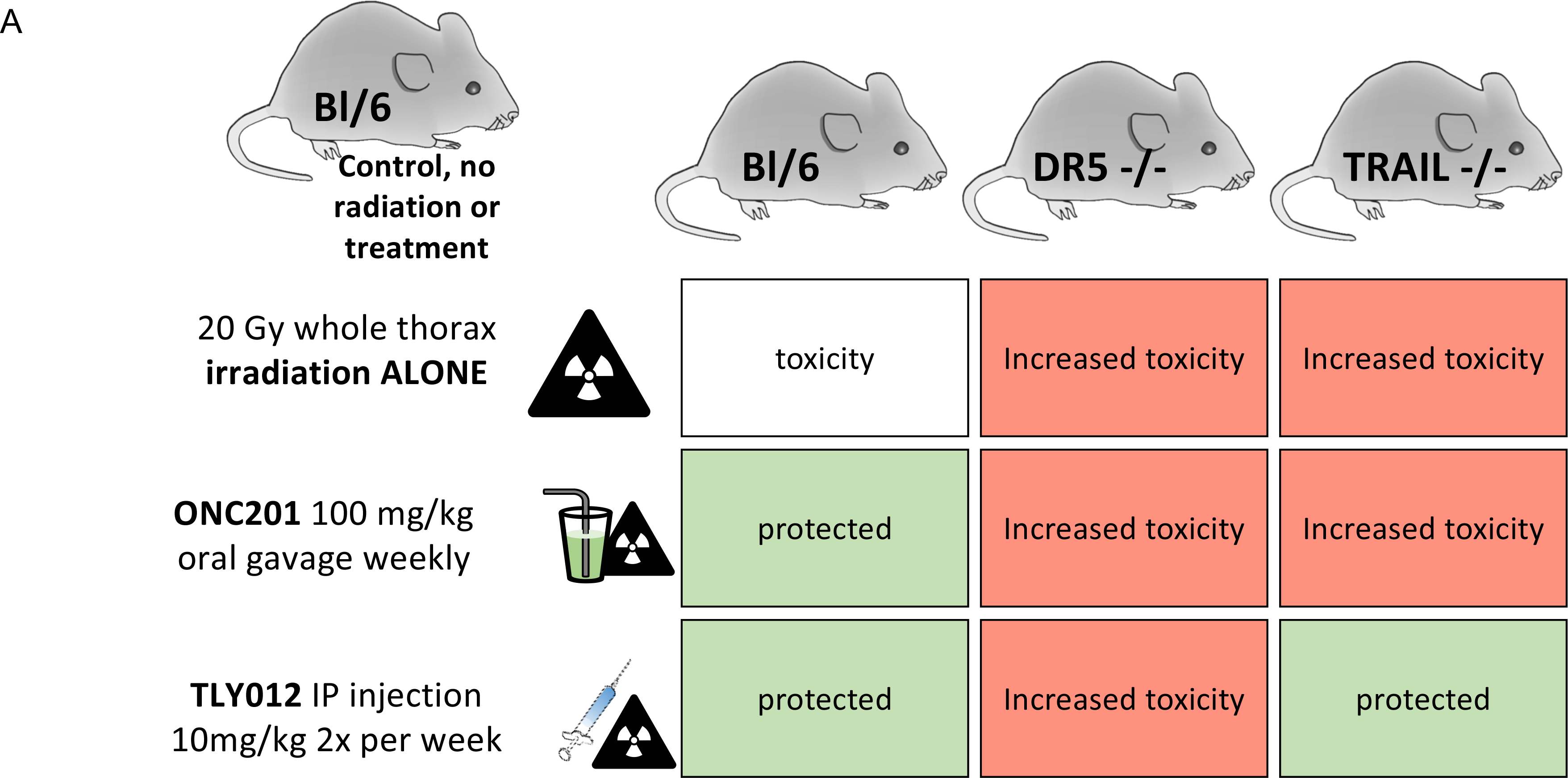
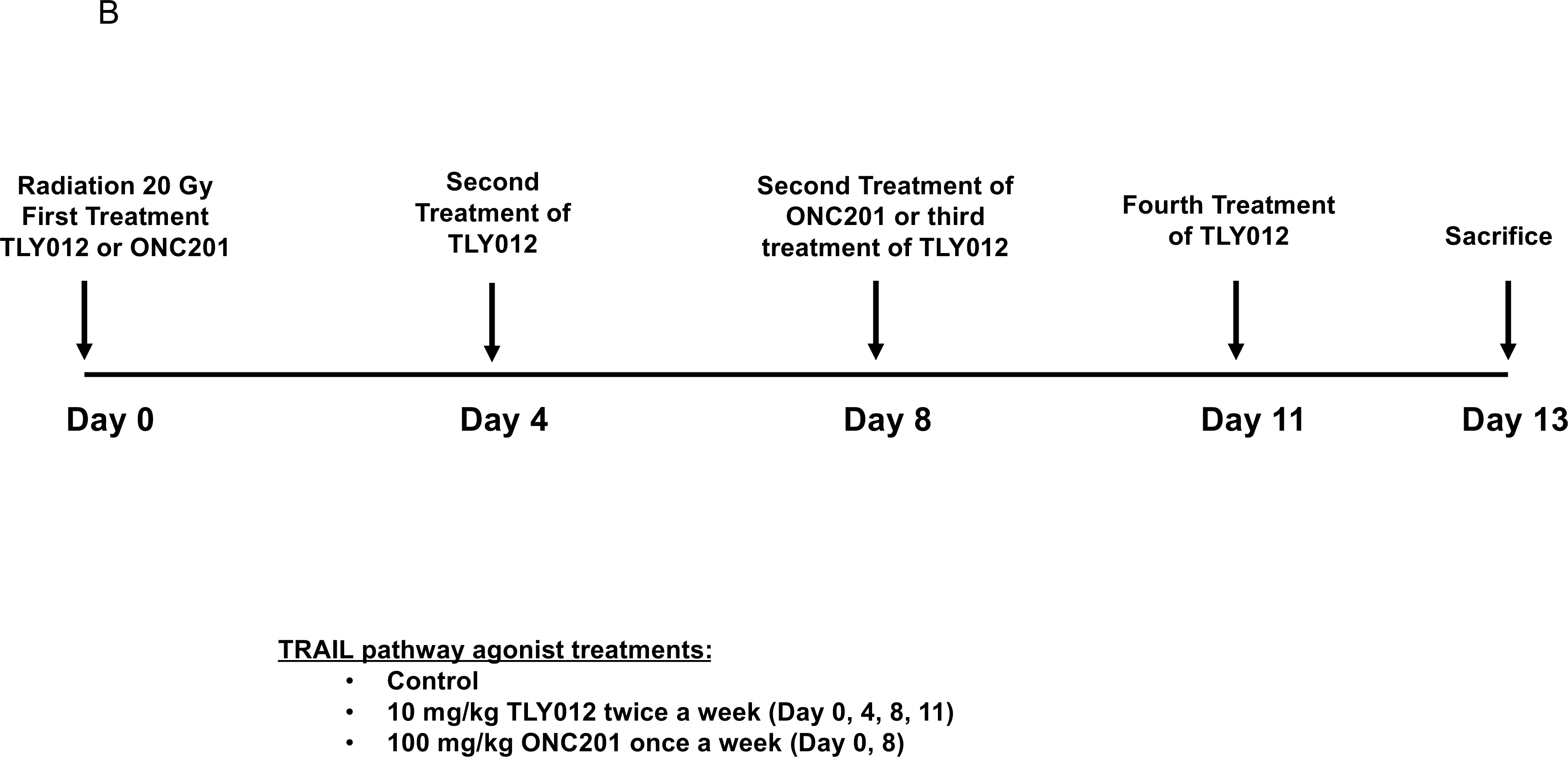
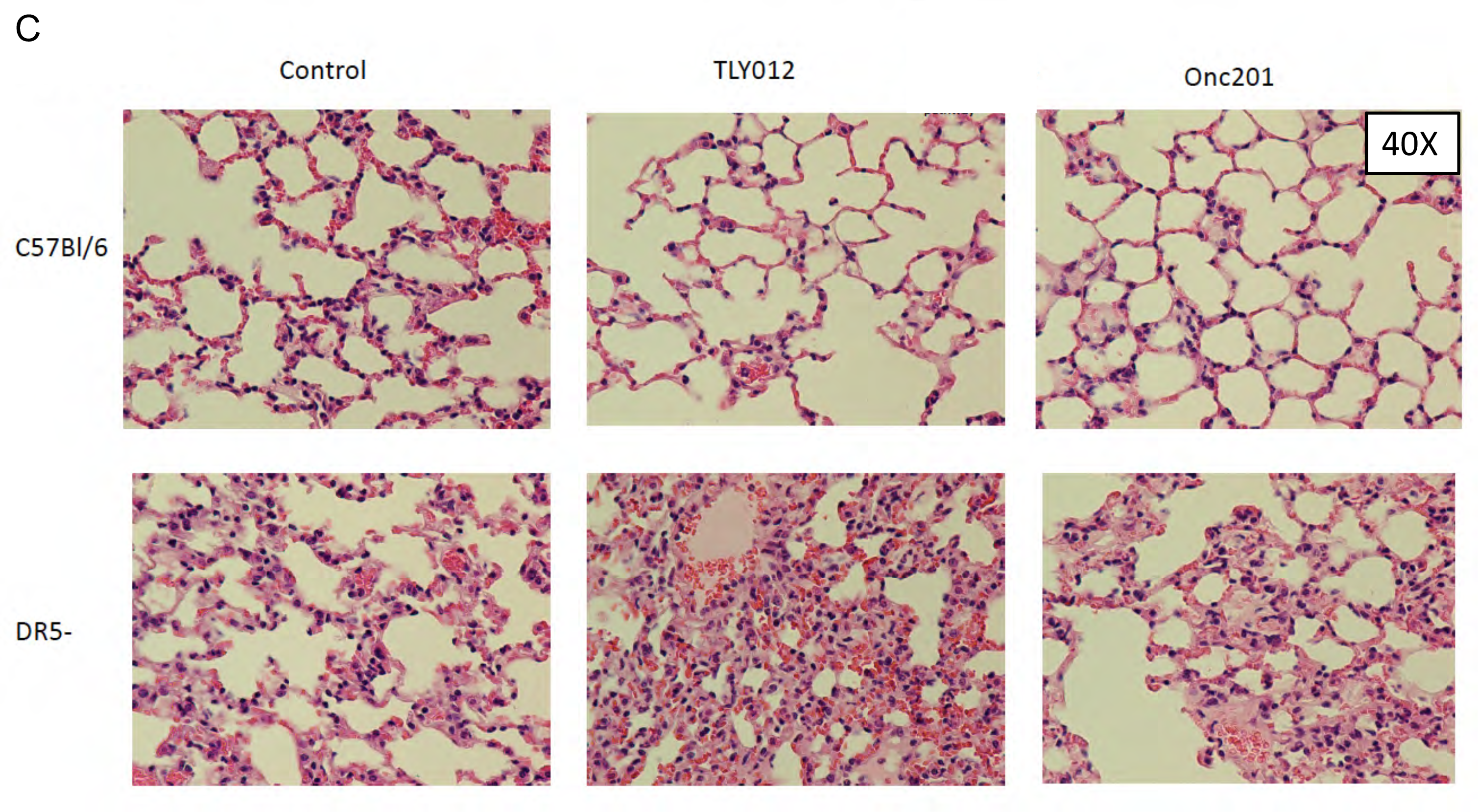
Experimental design including mouse strains and treatment cohorts A) Hypothesized outcomes regarding lung protection of C57Bl/6, *DR5*-/-, and *TRAIL*-/- mice irradiated with a whole-thorax x-ray irradiation dose of 20 Gy and treatment of either 100 mg/kg of ONC201 weekly, 10 mg/kg of TLY012 twice a week, or no treatment. **B)** Preliminary experimental timeline where mice received their first treatment just before radiation, and then were treated with either ONC201 once a week or TLY012 twice a week and then sacrificed on day 13 post-irradiation. **C)** First observations of suppression of radiation pneumonitis seen in x20 H&E-stained lung tissue of male mice by short term treatment with TRAIL pathway agonists as indicated (n = 2/treatment/genotype).

We performed a preliminary, short-term treatment feasibility experiment to ensure some mouse survival after 20 Gy of thoracic irradiation (**Figure 1B**). The preliminary experiment, performed with WT and *DR5*-/- mice irradiated with 20 Gy, was followed by either weekly doses of TIC10/ONC201 (100 mg/kg for total of two doses) or twice a week doses of TLY012 (10 mg/kg for total of four doses) for 2 weeks. Unexpectedly given the brevity of attempted rescue treatment, as shown in **Figure 1C**, we observed qualitative evidence of protection from pneumonitis in WT mice treated with either TLY012 or ONC201 versus control (20 Gy-irradiated mice that did not receive either TLY012 or ONC201). Lung inflammation in irradiated-mice was significantly worse in DR5-/- mice regardless of whether they were treated with TLY012 or ONC201 (**Figure 1C**). It was surprising that few doses of either TRAIL pathway agonist appeared effective in protecting against the severe radiation-induced acute lung injury. The lack of protection of DR5-/- mice and the appearance of a more severe phenotype is consistent with our previous observations (*17*). No obvious toxicities were noted in duodenum, liver or heart (**Figure S1 A-C**). Toxicity was also monitored by weighing the mice twice a week (**Figure S1 (D-F).** In this experiment, the lungs were not reinflated post-mortem, but it was determined that there were no significant visual differences between inflated versus un-inflated lungs (**Figure S1 G).**

### Rescue of WT or *TRAIL*-/- mice but not *DR5*-/- mice from radiation pneumonitis by TLY012

We tested the prediction that *TRAIL*-/- mice would be rescued by TRAIL (TLY012) but not by TIC10/ONC201 (**Figure 1A**). As predicted, short-term treatment with TLY012 over two weeks rescued 20 Gy irradiated (thoracic irradiation) male (**Figure 2A**) or female (**Figure 2B**) mice while ONC201 did not. This is consistent with the notion that if the TRAIL gene is not present, then ONC201 would not be expected to increase its expression of the gene and rescue pneumonitis. The results in **Figure 2A, B** demonstrate for the first time not only the severe lung inflammation in 20 Gy-irradiated *TRAIL*-/- mice, but also their rescue by TRAIL formulation TLY012. As hypothesized, we did not observe rescue of *DR5*-/- 20 Gy-irradiated lungs by either TLY012 or ONC201 (**Figure 2A, B**). We note, unexpectedly, that the severity of radiation-induced pneumonitis was worse in female versus male mice. Prior studies have either noted more toxicity in male rats or patients (*23–25*). In the case of the patients, it may be that more men had lung cancer and were treated with radiation.

**Figure 2.**
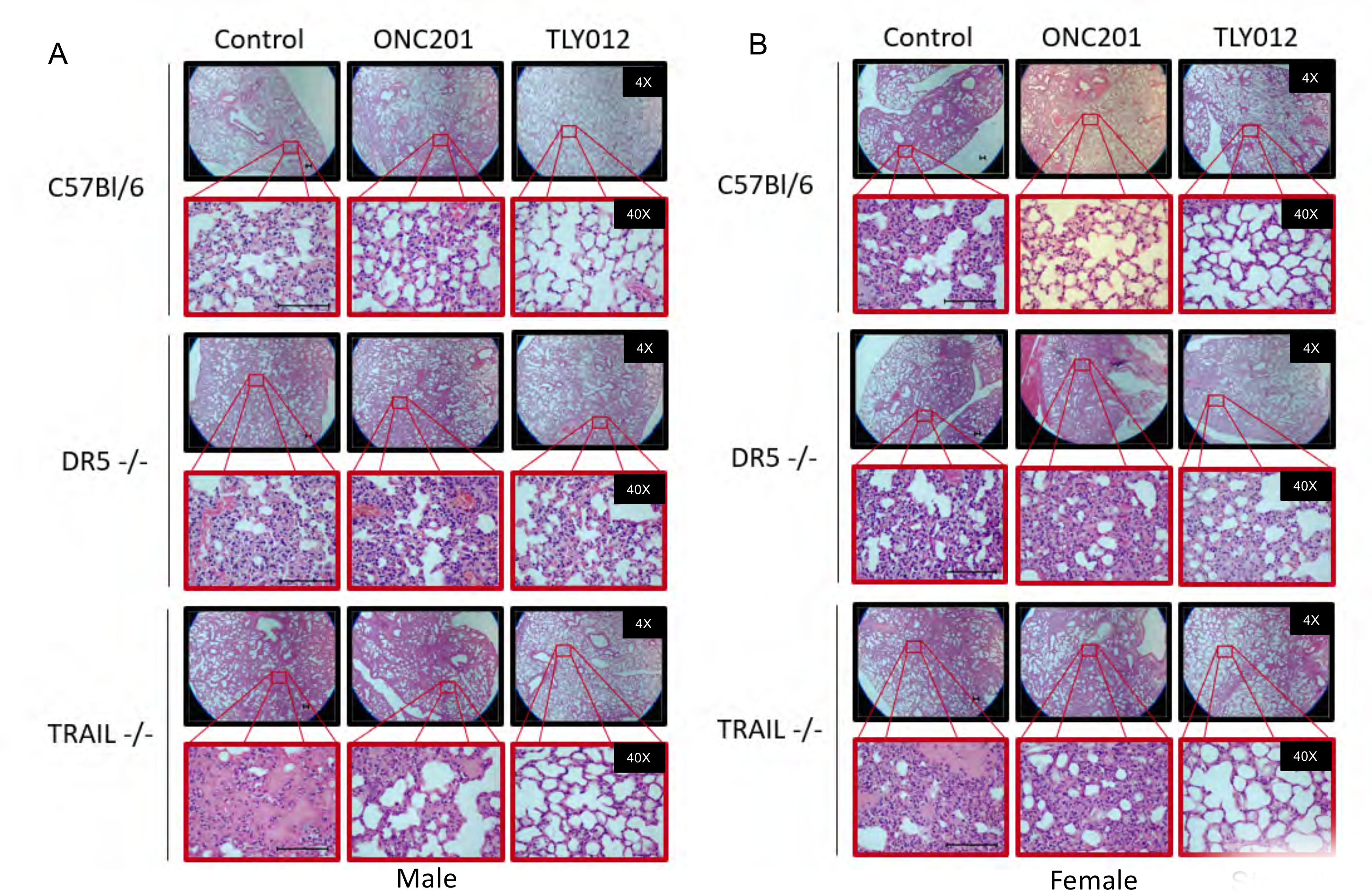
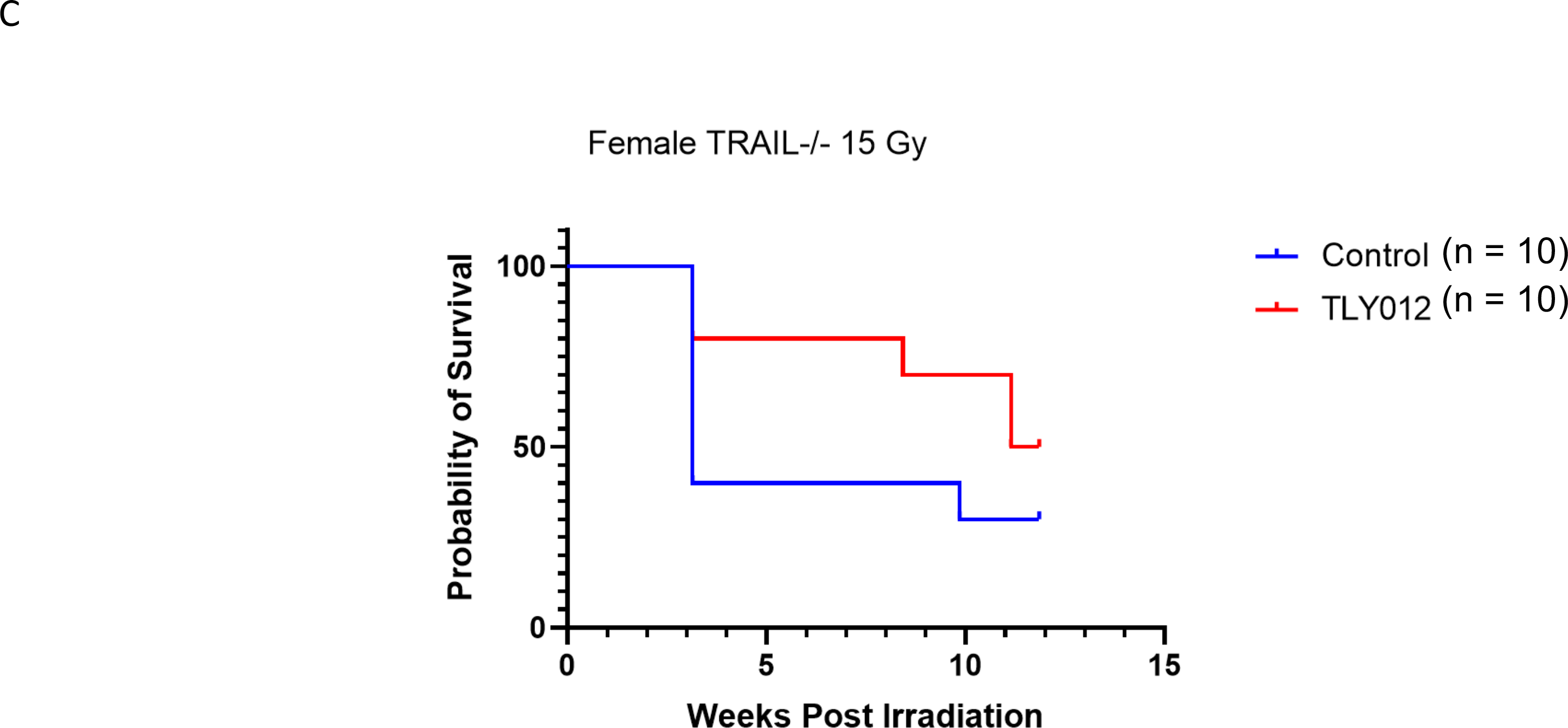
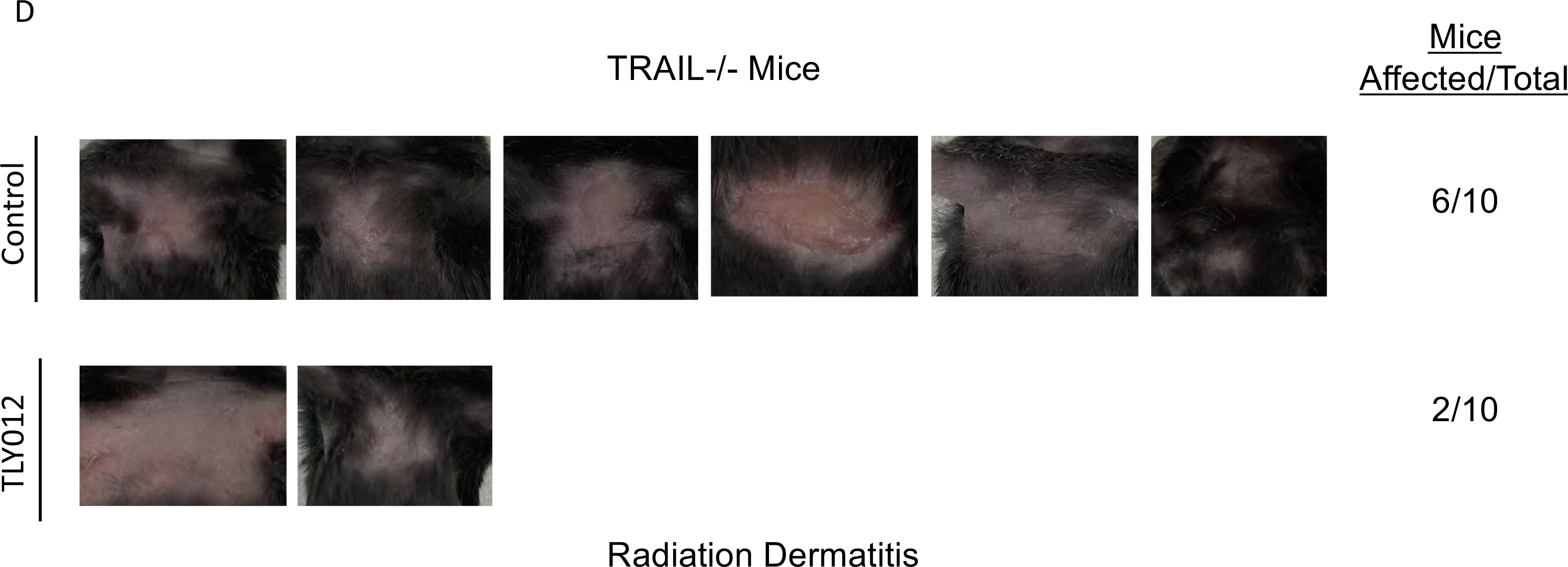
Protection of mice from radiation pneumonitis and dermatitis depends on genetic strain and TRAIL pathway agonist used to investigate mechanism of radioprotection. A,B) H&E stains of lung tissue from male and female mice from C57Bl/6, *DR5*-/-, and *TRAIL*-/- backgrounds treated with either ONC201, TLY012, or control gavage (n = 2/gender/genotype/treatment) 13 days post-irradiation (top row x4, bottom row x40) scale bar: 100 μm. **C)** Kaplan-Meier curve of TRAIL-/- female mice rescued from radiation dermatitis after a single whole-thorax x-ray irradiation dose of 15 Gy treated with TLY012 compared to control group (n = 10/treatment/group). **D)** Mice that developed radiation burns 3 weeks post irradiation on the chest were euthanized (control n = 6, TLY012 n = 2).

### Rescue from radiation dermatitis in TLY012-treated *TRAIL*-/- mice

Female *TRAIL*-/- mice that were given a single whole-thorax x-ray dose of 15 Gy (n = 10/treatment/group) developed radiation dermatitis in the chest area at ∼3 weeks post-irradiation (**Figure 2C)**. These burns evolved rapidly over 24-48 hours once they started to develop and the affected mice were euthanized per IACUC protocol. This unforeseen radiation toxicity was observed in *TRAIL*-/- mice and not in WT mice at the doses used. Of the *TRAIL*-/- mice that developed radiation dermatitis, 6 were control mice and 2 were TLY012-treated mice (**Figure 2D)**. This indicates that TLY012 is providing a protective effect against the radiation-induced skin damage as fewer TLY012-treated mice displayed severe rapidly-evolving dermatitis in the preliminary experiment. In addition to protection from radiation dermatitis, it is noted that up to 13 weeks post-irradiation, mice treated with TLY012 have a greater survival rate compared to control-irradiated mice (**Figure 2C).**

### Cytokine alterations in TLY012-treated mice

We examined patterns of cytokines in WT C57Bl/6, *DR5*-/- and *TRAIL*-/- mice that were irradiated and either not subsequently treated (control) or that were treated with TLY012 or ONC201 (**Figure S2 A, B**). We observed that IGF1 was upregulated by ONC201 while TLY012 increased CXCL1, IL6, GDF-15, IL6, MMP-8, and CCL3/MIP-1alpha (**Figure S2 A, B**). An increase in FGF-basic was noted with either TLY012 or ONC201. Prior work has implicated a range of other cytokines including TGF-beta, IL6, TNF and other inflammatory cytokines (*26*). We believe our results are providing initial insights for factors whose altered secretion patterns is evident by the effective pneumonitis-preventing agents that target activation of the TRAIL pathway.

### TLY012 protects from lethal radiation pneumonitis in *TRAIL*-/- mice

We investigated whether TLY012 could rescue mice from the lethality of thoracic irradiation (**Figure 3**). We observed rescue from lethality of *TRAIL*-/- male C57Bl/6 mice with 10 mg/kg dosing with TLY012 after a dose of 18 Gy of chest irradiation (**Figure 3A**). For female *TRAIL*-/- C57Bl/6 mice we had to reduce the radiation dose to 15 Gy to observe significant protection from lethality (**Figure 3B**). There was no protection from lethality by TLY012 of WT female C57Bl/6 mice after a high single dose of 25 Gy (**Figure 3C**).

**Figure 3.**
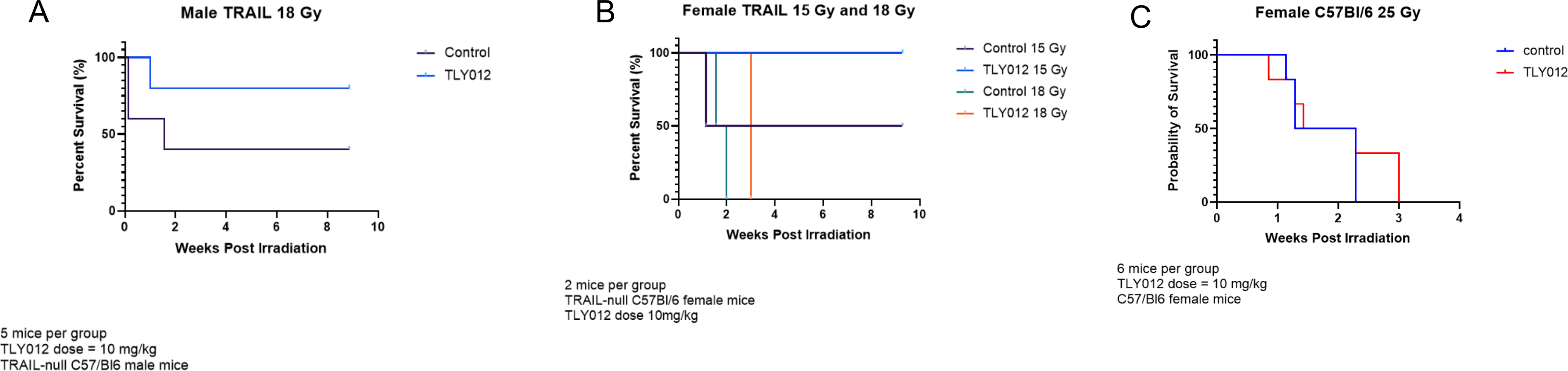
Protection for chest-irradiation induced mouse lethality by TLY012. A) Survival of male *TRAIL*-/- mice following 18 Gy irradiation led to an increased survival in mice treated with 10 mg/kg of TLY012 at 10 weeks following whole-thorax irradiation (n = 5/treatment/group). **B)** Survival of female *TRAIL*-/- mice following 18 Gy whole-thorax irradiation was increased to 3 weeks in mice treated with TLY012 compared to control mice that survived 2 weeks after radiation (n = 2/treatment/group). **C)** Survival of female C57Bl/6 mice was increased by approximately a week in mice treated with TLY012 compared to control mice (n = 6/treatment/group).

**Figure 4.**
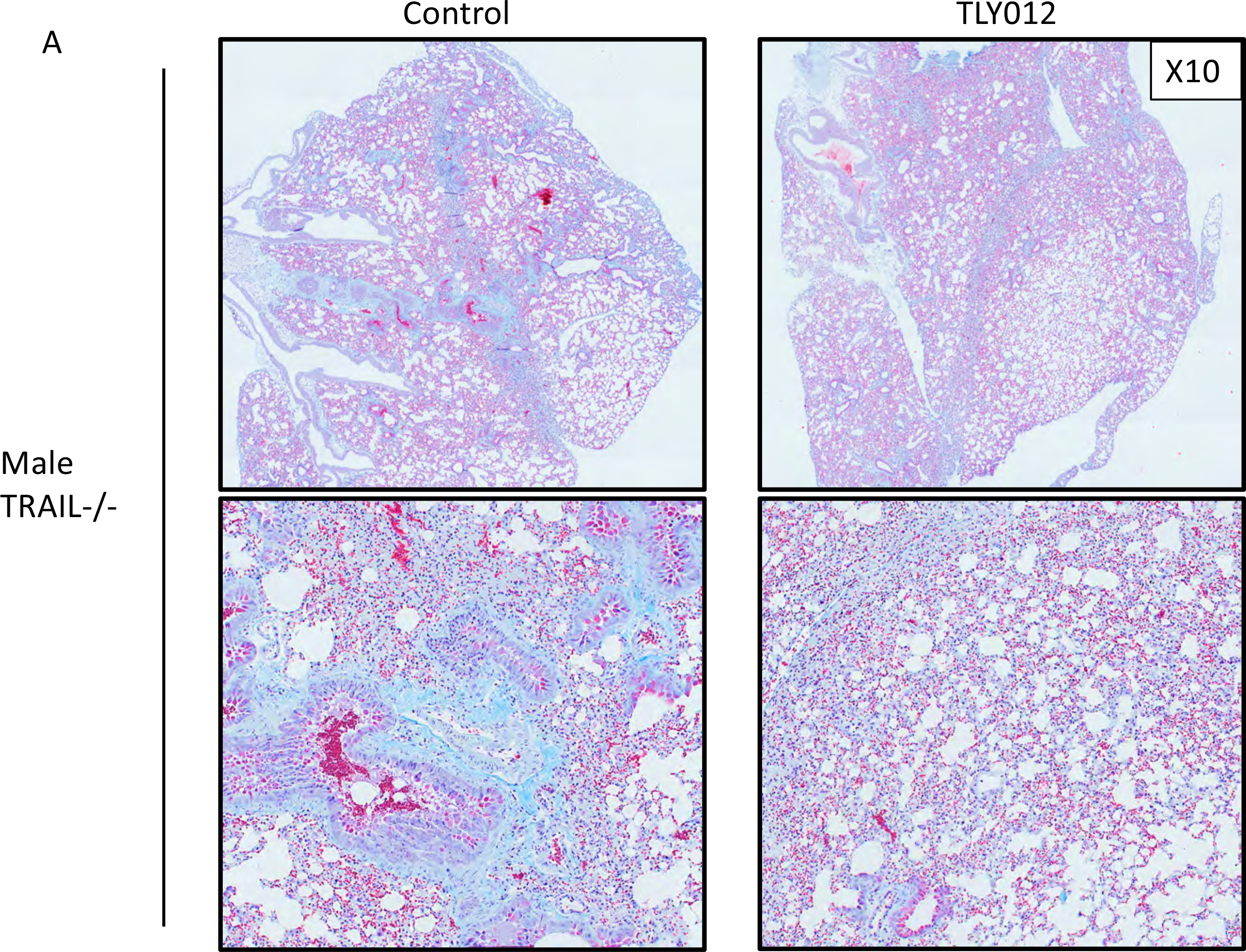
Masson’s Trichrome stain of male *TRAIL*-/- mouse lung 22 weeks post-radiation show rescue in TLY012 treated mice. A) Male *TRAIL*-/- mice that were treated with a single thoracic x-ray irradiation dose of 18 Gy and were treated with TLY012 or remained as control for 22 weeks post-radiation (n = 3, n = 2). Lung tissue was stained with Masson’s Trichrome and imaged at X10. (red = muscle fibers, bright blue = collagen, dark red/blue = nuclei).

### Reduced inflammation, and DNA damage in lung-irradiated versus TLY012-rescued mice

Examination of lung tissue in irradiated C57Bl/6 mice revealed that while TLY012 or ONC201 could rescue mice from radiation-induced pneumonitis (**Figure 2**), there is no difference in T-cell infiltration as judged by the pan T-cell marker CD3-epsilon (**Figure S3A**) or the DNA double strand break marker γ-H2AX (**Figure S3B**) within what remained as normal lung or what was inflamed lung with pneumonitis. It is clear, however, overall there are fewer areas of pneumonitis in rescued mice (**Figure 2**) but no difference in the marker expression in the analyzed healthy tissue or inflamed tissue in control versus treated mice (**Figure S3A, B**). Thus, the difference is the amount of inflammation in irradiated versus rescued mice rather than differences in the markers tested between normal appearing or injured areas of the lungs. Similarly, there are no differences in NFkB or cell proliferation between irradiated controls and irradiated rescued lung tissues (**Figure S3C, D**), although there is much greater overall areas of acute inflammation in irradiated versus TRAIL pathway agonist-rescued mice. There is less collagen deposition in rescued lungs (**Figure S3E**). Thus overall, there is less inflammation, DNA damage and collagen deposition in rescued lungs although in any residual inflamed tissues there is no difference in those regions between irradiated and TRAIL pathway agonist-rescued mice. While p53 expression appears similar across treatment groups, there is an observed increase in p53 expression in female mice of all three genotypes following irradiation (**Figure S3 F-H).**

### Decreased collagen production in mice after 18 Gy thoracic radiation and TLY012

To examine the late-term effects of thoracic radiation, we designed an experiment where the single radiation dose was lowered to 18 Gy in male *TRAIL*-/- mice. For 22 weeks post-radiation, mice were either treated with 10 mg/kg of TLY012 twice a week or remained control. At 22 weeks when mice reached criteria for euthanasia, they were sacrificed and lung tissue was harvested. Lung tissue slides were stained with Masson’s trichrome and imaged at 10X on an Olympus VS200 slide scanner. Upon analysis, there is significantly more light blue staining seen in control irradiated-mice as compared to TLY012-treated mice. The decreased collagen deposition in TLY012-treated mice suggests that a decrease in inflammation and rescue from late-term effects of radiation pneumonitis.

### Altered mRNA levels of genes related to inflammation and immune response

To determine the change in immune response between mice treated with TLY012 and controls after a single dose of 20 Gy thoracic x-ray irradiation, a NanoString PanCancer Immune Profiling panel was used to analyze mRNA from mouse lung tissue. It was found that 16 genes have statistically significant differential expression between the TLY012-treated group and the control radiation only group. *TNFSF10, KLRA7, CCL6, TMEM173, RELB, HERC6,* and *IL1RL2* were among the top upregulated differentially expressed genes (DEGs) in the TLY012-treated group relative to the radiation only control group, while *DOCK9, MAPK8, H2-Q2, PTGS2, RAET1A, BCL6, FOXJ1, IKZF2,* and *RRAD* are among the top downregulated DEGs (**Fig. 5A, B)**. NanoString pathway changes between the TLY012 treatment group and the control group were also examined. There are significant increases in “Antigen Processing” and “Interferon” pathways (**Fig. S5 A-D)**. When treatment TLY012 treated group and control group were further separated into male and female cohorts, it was found that there are significant increases in the “Antigen Processing”, “MHC”, and “Dendritic Cell Functions” pathways and a decrease in the “Basic Cell Functions” pathway in the female TLY012-treated mice.

**Figure 5.**
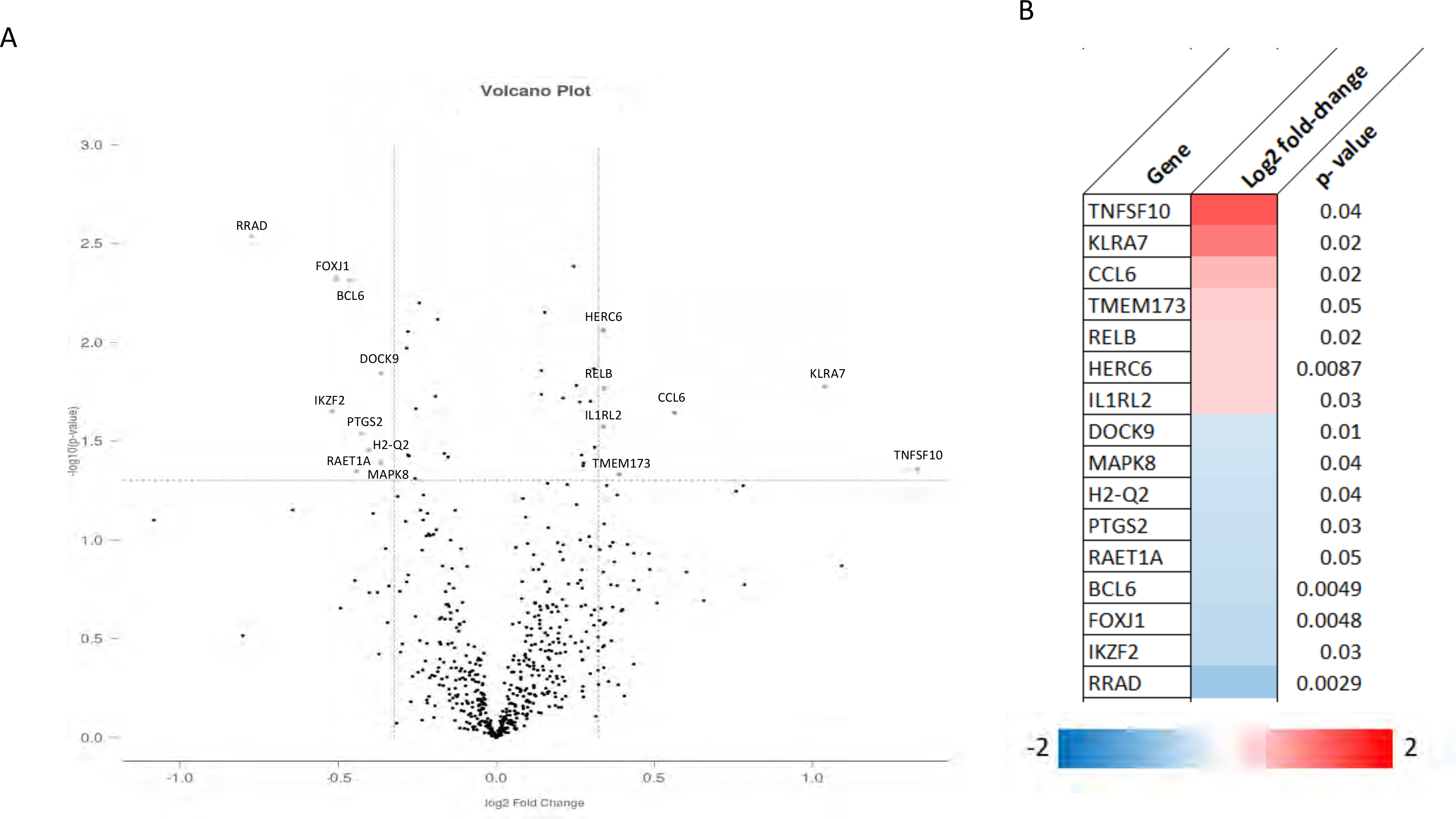
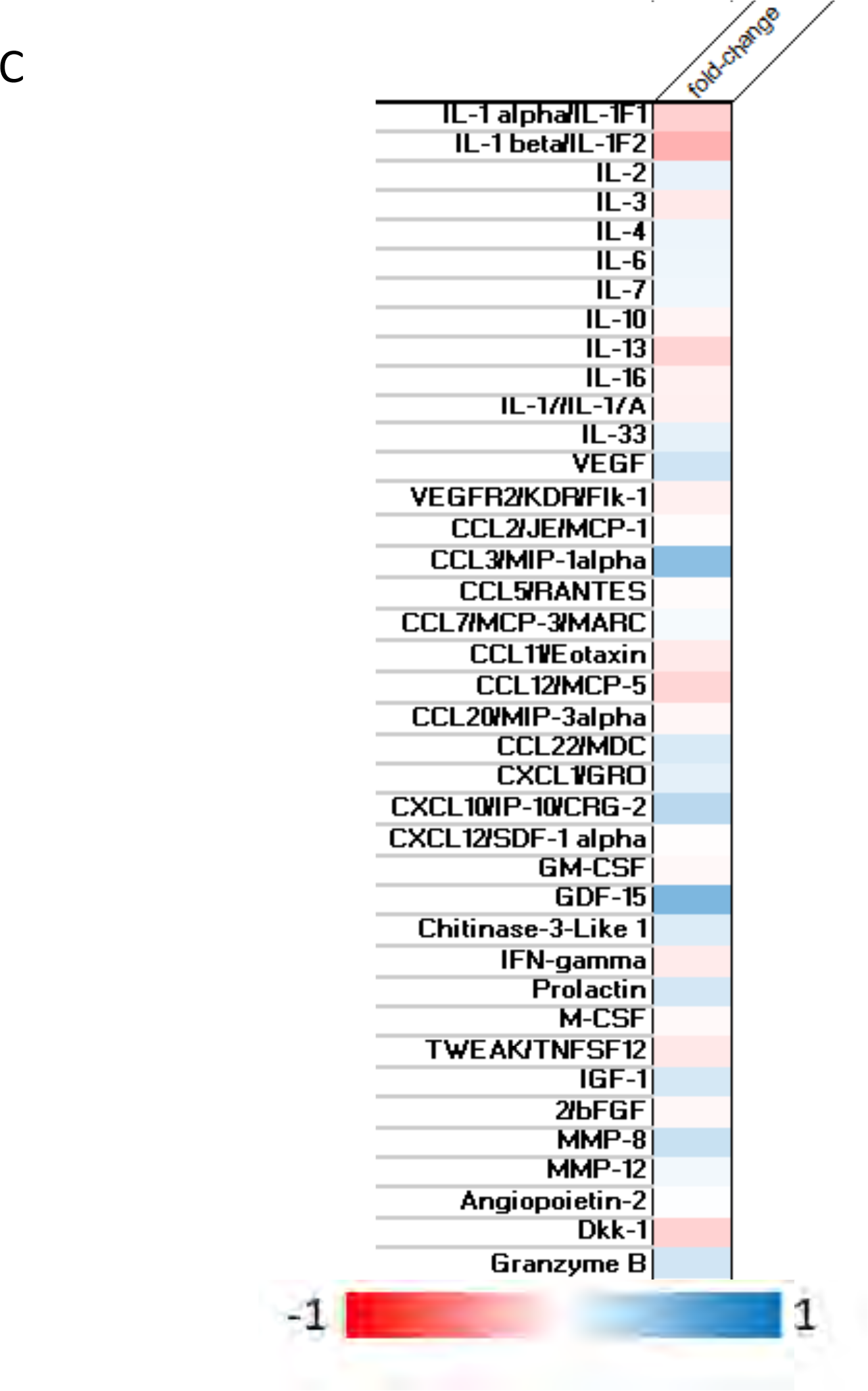
Nanostring nCounter PanCancer Immune panel for *TRAIL*-/- mice with and without rescue from radiation injury by TRAIL agonist. A) Volcano plot of all 40 references genes and their fold change compared to control mice. **B)** Heat map of log2 fold change of the top16 significantly different genes between control and TLY012 treatment (n=6). **C)** Heat map of serum cytokine level fold change of TLY012 treated mice relative to untreated control mice.

In addition to the NanoString PanCancer Immune profiling panel, serum cytokine analysis was performed. The cytokines that had the greatest differential expression in the TLY012-treated mice include a decrease in CCL3/MIP-1alpha, GDF-15 as well as an increase in IL-1beta (**Fig. 5C**).

### Orthotopic breast tumor-bearing immune-competent mice receiving 20 Gy thoracic radiation are protected from pneumonitis while showing complete elimination of tumors

We set up an experiment to determine whether TLY012, ONC201 or the combination used to prevent radiation pneumonitis might interfere with anti-tumor effects of radiation treatment to an orthotopically-implanted breast cancer in immune competent mice. We did not expect either ONC201, TLY012 or the combination to block anti-tumor efficacy given prior work (*27, 28*), but needed to formally show this in the radiation pneumonitis context. We injected mouse breast cancer e0771 cells orthotopically into mammary fat pad #2 of immune-competent C57Bl/6 mice and subsequently administered 20 Gy of radiation to the chest on day 9. Mice received TLY012, ONC201, the combination or no further treatment on days 9, 12, and 16 and then mice were sacrificed on day 18 (**Figure 6A**) as they ultimately began to lose more than 20% body weight in most treatment groups (**Figure S6 A)**. We observed an anti-tumor effect of radiation with no evidence that ONC201, TLY012 or their combination reduces efficacy of treatment (**Figure 6B, C**). Oxygen saturation is reduced in treated mice versus unirradiated mice and this is partially rescued by TLY012 or ONC201 versus irradiated mice that received no further treatment (**Figure 6D**). Rescue of mice from radiation induced pneumonitis appears to be more potent by TLY012 than with ONC201. The combination of TLY012 + ONC201, while effective, do not appear to improve the protection from pneumonitis under the experimental conditions (**Figure S6B**). Through cytokine analysis, there are differences between the control and treatment groups (**Figure S6 C).** There is reduction in CCL22/MDC levels in TLY012-treated irradiated tumor-bearing mice. CCL22 (**Figure 6E**) is a macrophage-derived chemokine previously associated with radiation pneumonitis and pulmonary fibrosis (*29*).

**Figure 6.**
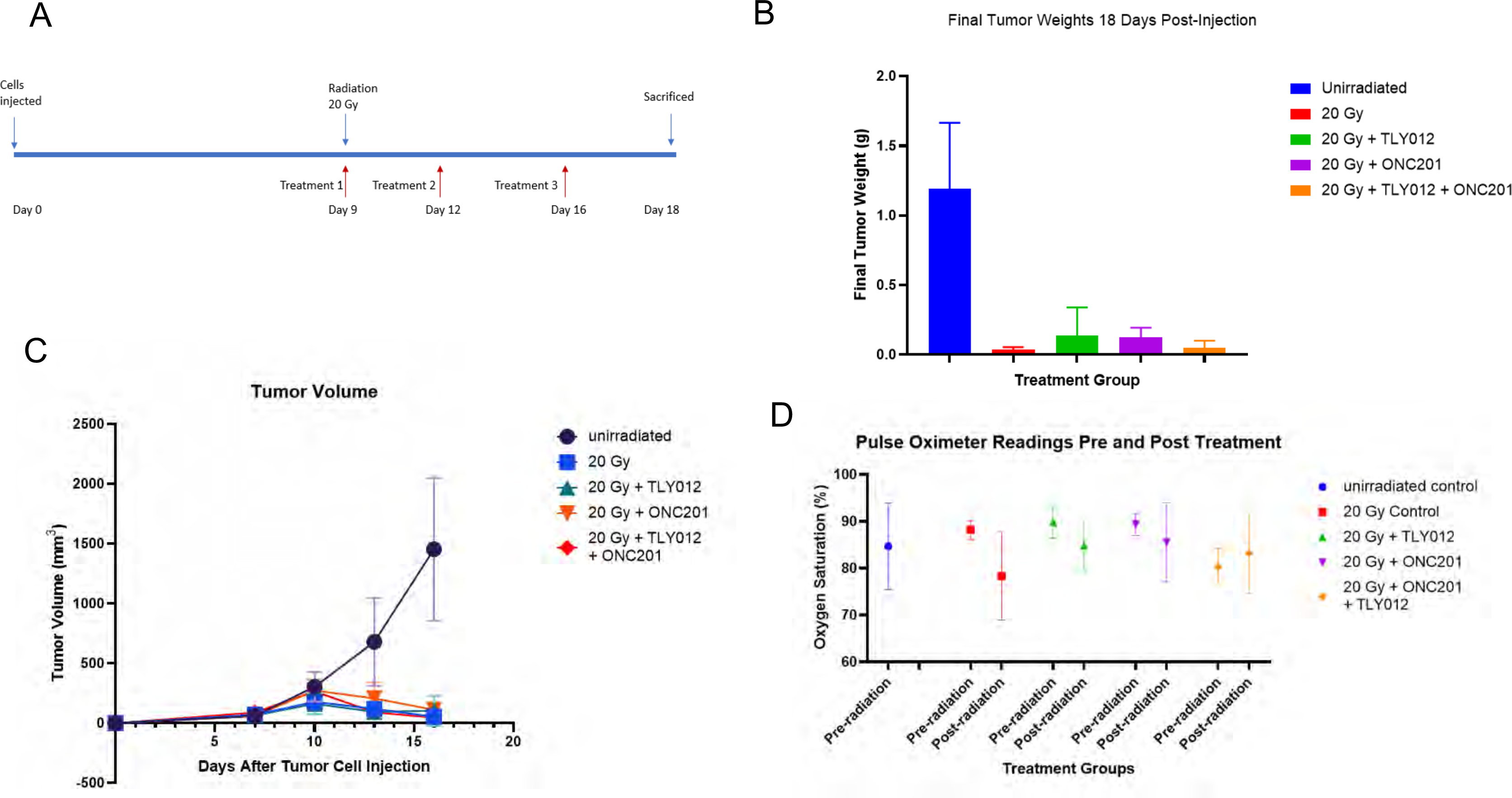
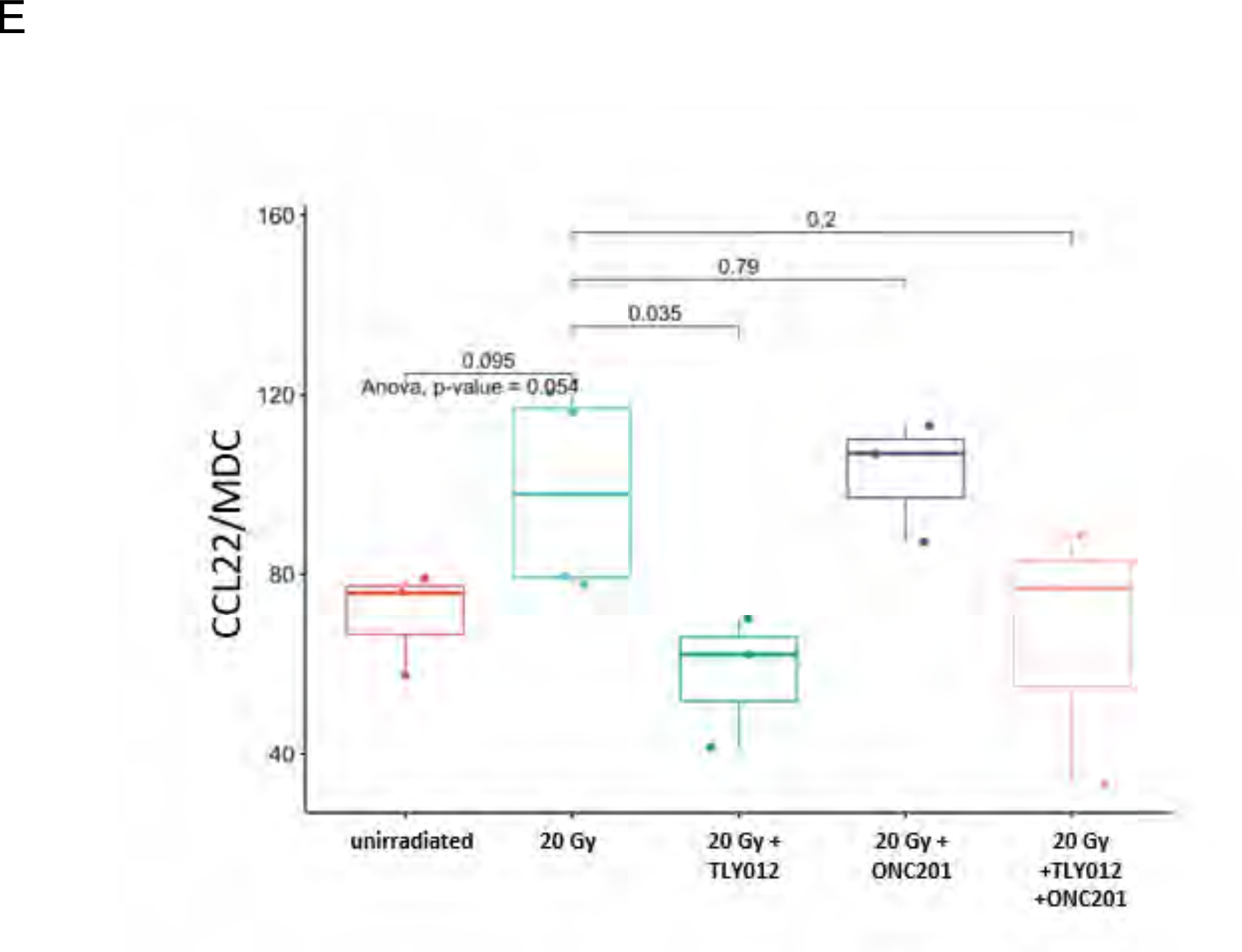
Efficacy of chest irradiation using an orthotopic immune-competent breast cancer model while preventing pneumonitis by TLY012 and reducing CCL22. A) Experimental timeline of female C57Bl/6 mice orthotopically injected with e0771 on day 0 and when tumors reached 2-5mm in size mice were irradiated with whole-thorax irradiation dose of 20 Gy and treated with either TLY012, ONC201, or a combination of both (n = 3/treatment/group). **B,C)** Tumors were removed after mice were euthanized 18 days post-cell injection where weight and volume were calculated. **D)** pulse oximetry readings pre-radiation and 9 days-post radiation showed oxygen saturation was more conserved in the TLY012 treated group compared to the irradiated control group. **E)** H&E stains of lung tissue 9 days-post irradiation showed a reduction in alveolar-wall thickness and decreased inflammation in mice treated with TLY012. **E)** Heat map of cytokine level fold change of treated mice to unirradiated control. **F)** Statistical analysis of cytokine fold change showed a significant decrease in levels of MDC/CCL22 (p = 0.035) when compared to the irradiated control group.

### *In vivo* micro-computed tomography (μCT) scans of mouse lung show rescue from toxic radiation-induced lung injury and fibrosis when treated with TLY012

In order to determine the effects of radiation on the mice *in vivo*, micro-computed tomography scans were taken of *TRAIL*-/- female mice that were unirradiated and two weeks-post one single thorax x-ray irradiation dose of 15 Gy on *TRAIL*-/- female mice with and without twice weekly TLY012 treatment (n = 2/treatment/group). Individual μCT image slices were collected as well as 3D reconstruction of the lungs during the inhale and exhale duration (**Figure S7 A)** of the breathing cycle. In the μCT mouse lungs expand more upon inhalation in the unirradiated and mice that were irradiated with 15 Gy and treated with TLY012 compared to the 15 Gy irradiated control. The distance between the heart and the esophagus is visually greater upon maximum inhale images in the TLY012-treated group as compared to the 15 Gy radiation only group, demonstrating the lungs are able to expand more **(Figure 7A).** In the 3D reconstruction (**Table S1)**, when all the images are subjected to the same opacity conditions, the unirradiated control and the 15 Gy irradiated group treated with TLY012 are overall clearer and have more visible airway space, particularly during the exhale portion of the breathing cycle **(Figure 7B)**. These observations are consistent with post-mortem findings through H&E staining and immunohistochemistry.

**Figure 7.**
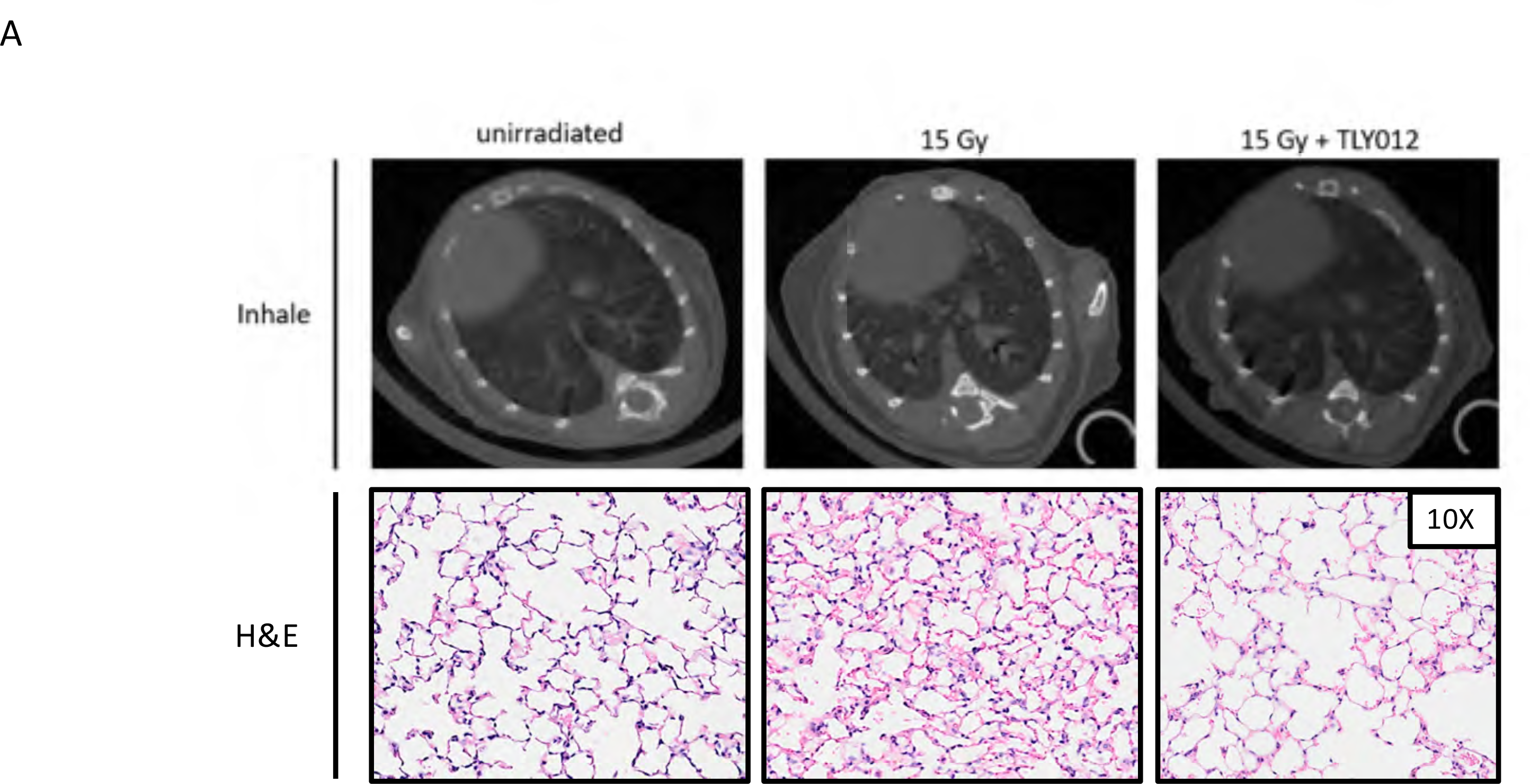
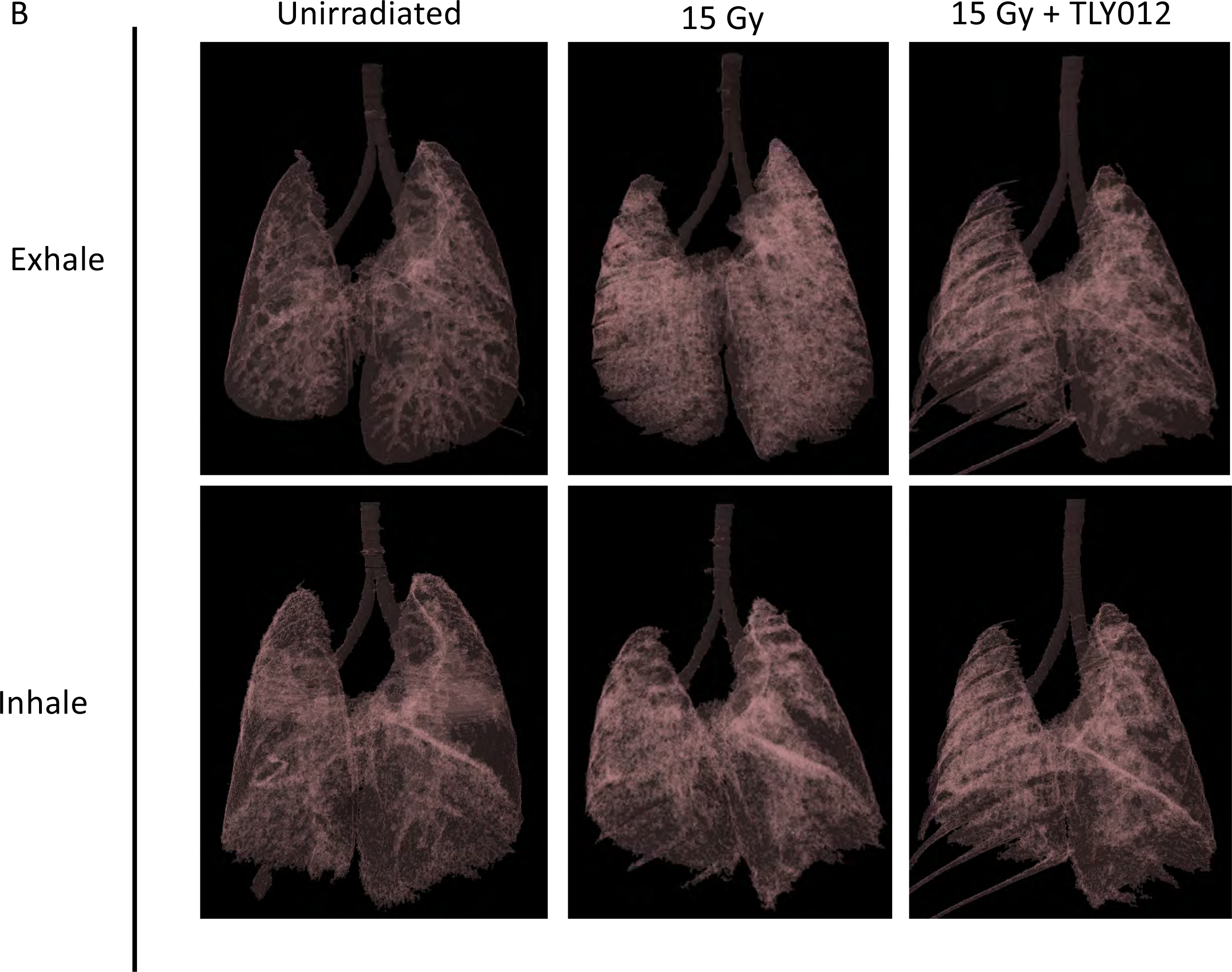
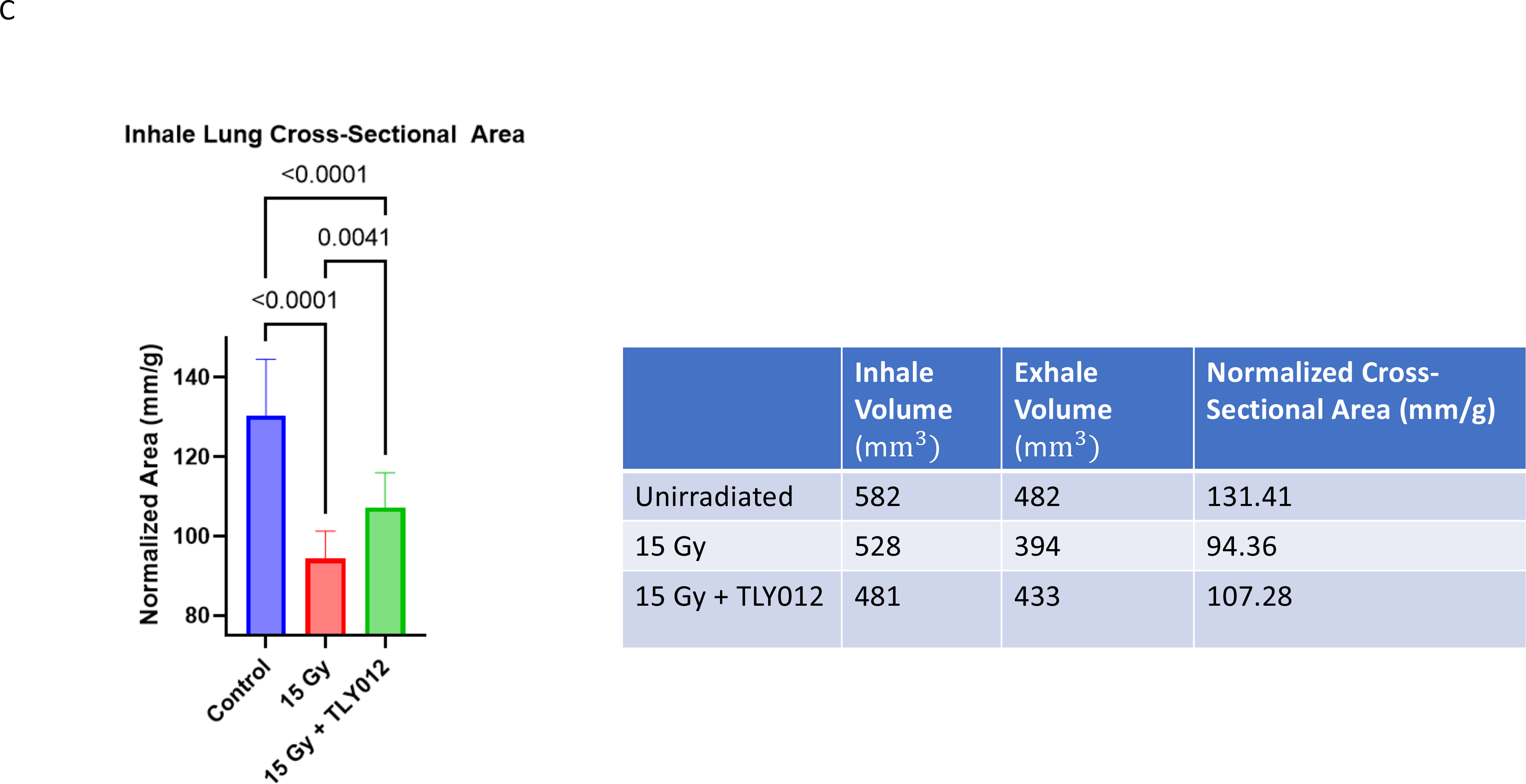
Respiration-gated imaging in mice irradiated with whole-thorax x-ray irradiation dose of 15 Gy with and without TLY012 treatment. A) Representative μCT images of mouse lungs were unirradiated, irradiated with 15 Gy, or irradiated with 15 Gy and rescued with TLY012 treatment during inhale duration of the breathing cycle (n=2/group). Mouse weights were 19.6 g, 24.1 g, and 22.4 g respectively. H&E slides of the lungs re-inflated post-mortem were also prepared. **B)** 3D reconstruction of mouse lungs from μCT images during exhale and inhale portions of the breathing cycle. All images are subjected to a 7% opacity filter in CTVol software. **C)** Area of the lungs was quantified from 15 CT images per mouse (n=1/treatment/group).

The same experiment was repeated in female *TRAIL*-/- mice where the mice were either unirradiated control, or received a single whole-thorax x-ray irradiation dose of 20 Gy and either remained as radiation only control or were treated with TLY012 twice a week for two weeks (n = 3/treatment/group). µCT image slices are consistent with the original experiment and show decreased lung volume upon the inhale portion of the breathing cycle with radiation alone versus unirradiated control or TLY012 rescue of irradiated mice (**Figure S7 B).** There are some abnormalities in the µCT images in the 20 Gy irradiated control mice around the heart that may represent atelectasis or opaque infiltration. Upon 3D reconstruction of the µCT images, it is of note that the 20 Gy control mice have lungs that are more congested with less visible airway space compared to the control (**Figure S7 C)**. Partial rescue is noted in mice irradiated with 20 Gy and treated with TLY012.

### Preliminary experiments with DR5 agonist and delayed treatment of TLY012 showed minimally but promising efficacy in rescue

To determine if TLY012 administration was time sensitive, we conducted an experiment where mice were administered TLY012 at 48-hours post radiation instead of 1-hour before treatment that was performed in other experiments. This experiment conducted in female C57Bl/6 mice irradiated with 20 Gy displayed that the time of dosing is important for extent of rescue from lung injury. A more complete rescue of the lungs is observed in mice given TLY012 1-hour before radiation compared to mice that receive the drug 48-hours post radiation (**Figure S8 A).** The mice were also weighed twice a week and both treated and control mice stabilize after day 8 (**Figure S8 B).** While the 48-hour post-radiation treatment does not show as much rescue as what is observed with same day TRAIL pathway rescue, we plan to further investigate the administration of TLY012 at 24 hours post-radiation in order to determine if there is greater rescue.

Another experiment utilized DR5 agonist (MD5-1) to investigate if degree of rescue to irradiated mice as compared to TLY012. In this experiment, male *TRAIL*-/- mice were irradiated with 20 Gy and then treated with MD5-1 twice a week or given an isotype control. While MD5-1 provides some rescue, there are many areas of inflamed tissue throughout the lungs. The results are similar to the IgG isotype control, implying that the drug did not provide significant rescue in the preliminary experiment without further optimization (**Figure S8 C).** Mice were weighed twice a week and both IgG isotype control and the MD5-1 treated groups recover on day 8 post-radiation (**Figure S8 D).** The greatest rescue is observed when TLY012 is administered to the mice 1-hour before radiation.

## DISCUSSION

The impact of this work is in prevention of lethality or other severe adverse consequences of therapeutic or unanticipated radiation injury in lungs such as pneumonitis and fibrosis by short-term treatment with innate immune TRAIL pathway agonists. Although we initially discovered the connection between the TRAIL death receptor signaling pathway and radiation pneumonitis nearly two decades ago (*30*), not until recently was it feasible to test whether this could be addressed through pharmacological interventions. It is surprising that short-term treatment with either TRAIL pathway agonist TLY012 or TIC10/ONC201 for two weeks prevents pneumonitis or lethality. This has implications for management of lung toxicity from radiation or as an approach for radiation counter-measures. Prevention of acute lung inflammation prevents delayed effects of radiation on the lungs and chronic lung injury.

One of the limitations of this study is that the lungs were not re-inflated post-mortem. We conducted a separate study comparing the H&E staining of inflated lungs versus non-inflated lungs of unirradiated mice and concluded that the alveolar border thickness was not greatly impacted by inflation (**Figure S2C**). Consistently across all of our models the lungs were not inflated but the lung inflammation and its rescue were obvious and functionally documented *in vivo*. Another limitation of this study is the number of mice used in each experiment. While each individual experiment only included a relatively small number of mice, we continued to observe that TLY012 significantly mitigated radiation pneumonitis across multiple experiments in male and female mice.

While there are statistically significant changes in lung tissue mRNA levels of genes related to immune response, there is no statistically significant fold-change in the cytokine levels of relative inflammatory markers in serum. The lack of significance in cytokines could be due to the fact that cytokine levels were measured from serum and not from bronchoalveolar lavage (BAL). The mRNA was extracted from lung tissue while cytokine levels were measured from peripheral blood serum levels. In future experiments BAL could be used to determine if differential cytokine activity is more prominent in the lungs. It should be noted that serum samples were taken 3 days after last administration of TLY012.

We note and emphasize that there was no difference in DNA damage (γ-H2AX) or T-cell inflammation (CD3-epsilon) in irradiated lungs in healthy or inflamed areas of the lungs regardless of whether they are rescued by TRAIL pathway activation or not. However, rescued mice have far fewer areas of pneumonitis, physiologically relevant increase in oxygen saturation, and evidence of rescued alveolar border thickness by histological examination of the lungs. The prevention of the pneumonitis by the TRAIL innate immune pathway agonists has no detrimental effect on the use of therapeutic radiation with regard to anti-tumor efficacy. Our results provide important clues as to alterations in cytokine biomarkers such as CCL22 or others that are impacted by TLY012.

While H&E stained images of the lungs show a reduced inflammatory phenotype in mice treated with TLY012, changes in gene and cytokine expression show decrease in inflammation as well. Genes related to a decrease in inflammatory response are increased in TLY012-treated mice. Killer cell lectin-like receptor 7 (KLRA7) is expressed by mature NK cells in mice (*31*). RelB, a member of the NF-kappaB/RelB family is known to act as a transcription suppressor in fibroblasts which limits the expression of pro-inflammatory mediators which have a significantly increased expression in TLY012-treated mice (*32*). Transmembrane protein 173 (TMEM173) encodes protein stimulator of interferon genes (STING), a key player in host defense against damaged cells. TMEM173 plays an important role in normal pulmonary function as it was found that loss-of-function TMEM173 alleles in humans lead to pulmonary fibrosis (*33*). HERC6 mediates ISGylation, which is involved in DNA repair and autophagy (*34, 35*). Interleukin 1 receptor like 2 (IL1RL2), which has an anti-inflammatory effect and can regulate macrophage function is also upregulated in mice treated with TLY012 (*36*). All of these findings suggest that treatment with TLY012 can prevent the inflammation and fibrotic effects caused by radiation.

Decreased expression of genes related to a proinflammatory response are also observed in mice treated with TLY012. Dedicator of cytokinesis 9 (DOCK9), which is thought to play a role in in immune disorders, shows a decrease in mRNA expression in mice treated with TLY012 (*37*). Mitogen-activated protein kinase 8 (MAPK8), also referred to as c-Jun amino terminal kinase (JNK) is activated in response to various cellular stresses and promotes production of pro-inflammatory and pro-fibrotic molecules is also significantly decreased in treated mice (*38*). Prostaglandin-endoperoxide synthase 2 (PTGS2), also known as COX2 is an enzyme that is responsible for causing chronic inflammation (*39*). Retinoic acid early transcript 1 alpha (RAET1A) functions as a stress-induced ligand for NKG2D receptor which is expressed on cytotoxic immune cells is downregulated in TLY012-treated mice, suggesting the control mice are in a greater state of stress (*40*). Another indicator of inflammation, B cell leukemia/lymphoma 6 (BCL6), is decreased in treated mice (*41*).

There remain open questions and research directions of interest. These include further mechanistic studies in the role of the cytokines and chemokines in the pathogenesis of pneumonitis or rescue from it. There are major questions about cross-talk between the TRAIL pathway and the TGF-β pathway, Gas/STING, and other cytokine responses previously linked to radiation pneumonitis. The findings in our manuscript may have relevance and could be examined in other lung injury models such as adult respiratory distress syndrome, idiopathic pulmonary fibrosis, immune checkpoint blockade pneumonitis, COVID-19 pneumonitis, or toxicities of other chemotherapy agents such as bleomycin. Combinations with other therapies such as steroids or TGF-β pathway inhibitors could be studied and more work can be done to understand differences between males and females in how they develop lung inflammation and fibrosis and why the phenotype in females is more severe in *DR5*-/- and *TRAIL*-/- mice. Further studies can investigate extent of protection to other organs including bone marrow, GI tract, or brain from toxic effects of a range of radiation doses. More investigation can be carried out to determine whether TRAIL pathway agonists can impact on various forms of dermatitis along with more detailed studies of underlying mechanisms. More work can be carried out to determine how late in time TRAIL pathway agonists could impact on protection from radiation post-exposure including in combination with other agents. Our preliminary results suggest some rescue is possible even at 48 hours after irradiation and treatment by TLY012 (**Figure S8**). Our findings have translational relevance by suggesting future clinical investigation of the TRAIL innate immune pathway in toxicity of radiation or other cancer therapeutics, other inflammatory lung or skin conditions and have implications for the field of radiation countermeasures.

## MATERIALS AND METHODS

### Bioassays

Animal studies were carried out at Brown University facilities with approval from Brown University IACUC. C57Bl/6 mice (Taconic), mice with bi-allelic loss of TRAIL Death Receptor-5 (*DR5*-/-) in a C57Bl/6 background, and mice with bi-allelic loss of TRAIL (*TRAIL* -/-) in a C57Bl/6 background were given a single whole-thorax x-ray (Philips RT250) irradiation dose of 20 Gy with shielding of other organs. 8–15-week-old male and female mice of each genotype were used. Starting one hour before irradiation, mice were treated with either 10 mg/kg of TLY012 (D&D Pharmatech) by intraperitoneal (IP) injection or 100 mg/kg of ONC201/TIC10 (Chimerix/Oncoceutics) by oral gavage (PO) or a control gavage consisting of 20% Cremophor® EL (Sigma #238470) and 70% Dulbecco’s phosphate buffered saline solution (Cytiva SH30264.02) per volume via PO (n = 2/gender/treatment). Treatment was continued for two weeks, where TLY012 was administered twice a week and ONC201 was administered once a week. For each genetic background, two mice received a control PO, two mice received TLY012 IP, and two mice received ONC201 PO. Mice were weighed twice a week, and 13 days post-irradiation all mice were euthanized. Organs were harvested from the mice and 600 µL of blood was collected via cardiac puncture for serum cytokine analysis. Organs were preserved in 10% formalin then embedded in paraffin and sectioned at a distance of **5 µm**. Hematoxylin and eosin (H&E) stained slides of the lungs, heart, liver, and duodenum were imaged at 4X and 40X magnifications on a Nikon Y-THM Multiview Main Teaching Unit microscope using a Diagnostic Instruments, Inc. model 18.2 color mosaic camera paired with SPOT Basic version 5.3.5 software.

Survival experiments were conducted using male *TRAIL*-/- mice treated with either TLY012 or control exposed to a whole-thorax radiation dose of 18 Gy (n = 5/treatment) as it was determined any dose above 18 Gy caused too great of radiation toxicity in order to allow long term survival (**Figure 3A**). *TRAIL*-/- female mice were also subjected to a survival study with a single whole-thorax radiation doses of 15 Gy or 18 Gy treated with the same regimen of TLY012 or control (n = 2/radiation dose/treatment) (**Figure 3B**). In order to determine the lethal dose of radiation for C57Bl/6 mice, female C57Bl/6 mice were treated with a single dose of whole-thorax irradiation of 25 Gy in a survival study where they were treated with either TLY012 or remained control (n = 6/treatment) (**Figure 3C**).

### Tumor Bearing Model

C57Bl/6 female mice (Taconic) were injected with 100,000 cells of murine breast cancer (e0771) in the right second mammary fat pad on day 0 of the experiment. The e0771 breast cancer cell line is classified as Luminal B, ERα-, ERβ+, PR+, and Her2+ (*42*). On day 9, tumors were greater than 4 mm in diameter and mice were grouped into control, treatment of 10 mg/kg of TLY012 twice a week, 100 mg/kg of ONC201 once a week, or a combination of both drugs. Each group (*n*=3) also received a single whole-thorax x-ray irradiation dose of 20 Gy with shielding of other organs. One group (*n*=3) received no treatment and no radiation. The long axis and short axis of the tumors were measured using calipers and recorded twice a week. On day 18 all mice were euthanized at a humane endpoint, which was defined by 20% weight loss and/or a tumor volume of 2000 mm^3^ (∼10% of body weight). Organs and tumors were harvested from the mice and 600 µL of blood was collected via cardiac puncture for serum cytokine analysis. Once tumors were extracted, they were weighed and measured to determine final volume and weight. Pulse oximeter readings were taken initially before radiation treatment and before sacrifice 9 days post-radiation (MouseSTAT Jr. Pulse Oximeter, Kent Scientific). When mice were under anesthesia the sensor of the MouseSTAT Jr. was placed on the hind right paw for 30 seconds. The highest and lowest values shown by the pulse oximeter during this time were recorded and then averaged together. The mean of all the averages per treatment group was then calculated.

### DR5 agonist

8–13-week-old male *TRAIL*-/- mice were given a single whole-thorax x-ray irradiation dose of 20 Gy with shielding of other organs. An hour before radiation, mice were treated with 100 μg anti- DR5 mAb (clone MD5-1; BioXCell) or given 100 μg isotype control of polyclonal Armenian hamster IgG (BE0091; BioXcell) via IP injection (n=3/treatment). Mice were treated once a week for two weeks and sacrificed on day 13-post irradiation. Lungs were harvested and preserved in 10% formalin then embedded in paraffin and sectioned at a distance of 5 µm. H&E-stained slides of the lungs were imaged at 20X magnifications.

### TLY012 treatment at 48-hour post-radiation

Female 8-week-old C57Bl/6 mice (Taconic) were given a single whole-thorax x-ray irradiation dose of 20 Gy with shielding of other organs. At 48-hours after administration of radiation, mice were grouped into control or treated with 10 mg/kg of TLY012 twice a week (n=4/treatment). Mice were weighed twice a week, and then euthanized on day 13 post-irradiation. Lungs were harvested and preserved in 10% formalin then embedded in paraffin and sectioned at a distance of 5 µm. H&E-stained slides of the lungs were imaged at 20X magnification.

### Cytokines

All mouse serum samples were analyzed by a custom R&D systems Murine Premixed Multi-Analyte Kit (R&D Systems, Inc., Minneapolis, MN). The panel was run on a Luminex 200 Instrument (Luminex Corporation, Austin, TX) according to the manufacturer’s instructions. Murine serum levels of Angiopoietin-2, BAFF/BLyS/TNFSF13B, CCL2/JE/MCP-1, CCL3/MIP-1 alpha, CCL4/MIP-1 beta, CCL5/RANTES, CCL7/MCP-3/MARC, CCL11/Eotaxin, CCL12/MCP-5, CCL20/MIP-3 alpha, CCL21/6Ckine, CCL22/MDC, Chitinase 3-like 1, CXCL1/GRO alpha/KC/CINC-1, CXCL10/IP-10/CRG-2, CXCL12/SDF-1 alpha, Dkk-1, FGF basic/FGF2/bFGF, GDF-15, GM-CSF, Granzyme B, IFN-gamma, IGF-I/IGF-1, IL-1 alpha/IL-1F1, IL-1 beta/IL-1F2, IL-2, IL-3, IL-4, IL-6, IL-7, IL-10, IL-13, IL-16, IL-17/IL-17A, IL-27, IL-33 M-CSF, MMP-3, MMP-8, MMP-12, Prolactin, TWEAK/TNFSF12, VEGF, and VEGFR2/KDR/Flk-1 were measured. Analyte values were reported in picograms per milliliter (pg/mL). TGF-β1 samples were run using a 1:15 dilution factor, as suggested by the manufacturer. Latent TGF-β1 was activated in samples using 1 N HCl and samples were neutralized with 1.2 N NaOH/0.5 M HEPES. Samples were assayed immediately after neutralization. Statistical analysis was performed using R software where one way ANOVA and post hoc Turkey test for pairwise comparisons were utilized.

### Immunohistochemistry (IHC)

Formalin-fixed paraffin-embedded sections of tissue were deparaffinized in xylene and rehydrated through graded ethanol solutions to phosphate-buffered saline (PBS). Heat-induced antigen retrieval (in 0.01 M citrate buffer; pH 6.0), endogenous peroxidase blocking (3% H2O2), and blocking were done using a standard protocol. Diluted primary antibody (CST #8242 1:800; CST #99940, 1:150; CST #9718, 1:500; Abcam ab107099, 1:X; Leica P53-CM5P-L 1:200) was applied and slides were incubated at 4°C overnight. Slides were then washed with PBS and incubated at room temperature for one hour with appropriate secondary antibodies (Vector Laboratories MP- 7401, MP-7402). Slides were developed with diaminobenzidine substrate (Vector Laboratories SK-4100) and counterstained with hematoxylin (Richard Allen Scientific). Slides were scanned (Olympus VS200) and representative brightfield images were taken and analyzed with OlyVIA software.

### NanoString Assay

Male and female 9-week old *TRAIL*-/- mice were given a single whole-thorax x-ray irradiation dose of 20 Gy with shielding of other organs. One hour before radiation mice were grouped into control or given 10 mg/kg of TLY012 via IP injection (n = 3/gender/treatment). Mice were treated and weighed twice a week up until 11 days post-radiation when all mice were euthanized.

Rodent sacrifice was performed following anesthetization with 100 mg/kg ketamine and 10 mg/kg xylazine administered intraperitoneally. ∼400-600 μL blood was first collected through cardiac puncture with a 26G 5/8” needle through the intercostal space into the left ventricle. Following cervical dislocation, the inferior vena cava (IVC) and descending abdominal aorta were severed prior to bilateral thoracotomy to expose the heart and lungs. 1 mL of 2 mg/mL EDTA in PBS was injected through the right ventricle to perfuse the lungs, after which the lungs were excised, washed further in PBS, submerged in RNAlater (Sigma-Aldrich, #R0901, Missouri, USA), and flash frozen in liquid nitrogen.

Frozen lung tissue was stored long-term at -80°C. Thawing was performed over 16 hours at 4°C. Tissue homogenization was performed with Precellys Lysing Kit (Cayman Chemical Company, #16859, Michigan, USA) and a Bulley Blender Homogenizer (Next Advance, #G14-G15) prior to RNA extraction with QIAgen RNeasy kit (Qiagen, #74104, Hilden, Germany) and cleaned up with QIAgen RNeasy MinElute Cleanup kit (Qiagen, #74204, Hilden, Germany). Quality and concentration were verified by Nanodrop. Extracted RNA samples were gene expression profiled by NanoString nCounter PanCancer Immune Profiling Panel (NanoString Technologies, #XT-CSO-MIP1-12, Seattle, WA) according to the manufacturer’s instructions.

Nanostring data was analyzed in nSolver Advanced Analysis Software and ROSALIND. Raw data was uploaded to nSolver for automated normalization, background subtraction, and quality control (QC) check. All samples passed QC. Control mice and mice treated with TLY012 lung samples were used to construct two groups to which an unpaired t-test was run to generate the data in the volcano plot that was created in ROSALIND (**Fig. 5)**. Differential expression was determined with p-values and Benjamini-Yekutieli adjusted p-values. Pathway scores are generated in nSolver as a summarization of expression level changes of biologically related groups of genes. Pathway scores are derived from the first Principal Components Analysis (PCA) scores (1st eigenvectors) for each sample based on the individual gene expression levels for all the measured genes within a specific pathway. The cell type score itself is calculated as the mean of the log2 expression levels for all the probes included in the final calculation for that specific cell type.

### *In vivo* microCT Imaging

For μCT imaging, mice were imaged in a SkyScan 1276 *in vivo* microCT scanner (Bruker, Kontich, Belgium) under isoflurane at 41 μm voxels with a 0.7° rotation step in supine position. Mice that underwent μCT of the lungs consisted of *TRAIL*-/- females that were unirradiated, and irradiated at 15 Gy with and without treatment of 10 mg/kg of TLY012 twice a week (n = 2/group). Mice underwent μCT scans at 13 days post-irradiation. Image-based respiratory gating was used to sort images into gates based on specific time points of the breathing cycle to minimize motion artifact in the final reconstruction (**Fig. 7A**). For 3D-reconstruction, Dataviewer version 1.5.4 (Bruker) was used to sort the CT images using the function “Listmode Scan” grouping the images in each bin where bin 0 = maximum exhale and bin 3 = maximum inhale (empty views <10%). Once the images were sorted, the listmode dataset was opened in NRecon software (Bruker) where the ROI undergoes 3D reconstruction. Reconstructed images were then analyzed in CTAn software (Bruker) where the volume of the lungs was calculated during max inhale and max exhale. The plugins in the custom processing page were used to create an ROI that separated the lungs from the rest of the image. 3D reconstructed images of the lung only were then opened in CTvol software (Bruker) where the color (red = 69%, green = 50%, blue = 50%) and the opacity (7%) of the lung was changed (**Fig. 7B**). H&E stained slides were also prepared post-mortem after lung reinflation with 1 mL of PBS and sectioning.

The area of the lungs was quantified via ImageJ utilizing the polygon function to separate the lungs from the rest of the body (**Fig. 7C)**. For each mouse, 15 different microCT images were measured and then averaged together (n = 1/treatment/group). As the microCT images were taken at 41 µm, the areas were converted from pixels to millimeters. The areas of the control mouse, 15 Gy irradiated mouse, and 15 Gy irradiated mouse treated with TLY012 were 131.41 mm/g, 94.36 mm/g, and 107.28 mm/g respectively. Statistical analysis was performed by utilizing GraphPad. The one-way ANOVA test showed the mouse treated with TLY012 had a significantly larger lung area compared to the irradiated mouse (p value = 0.0041).

The experiment was then repeated with female *TRAIL*-/- mice where mice were either unirradiated control, or received a single whole-thorax x-ray irradiation dose of 20 Gy and either received no treatment or TLY012 twice a week for two weeks (n=3/treatment/group). Two weeks post-irradiation, mice underwent µCT scans and then were euthanized. µCT scans underwent 3D- reconstruction for the inhale portion of the breathing cycle and were compared across treatment groups (**Figure S7 B-C).**

## Acknowledgements

W.S.E-D. is an American Cancer Society Research Professor and is supported by the Mencoff Family University Professorship at Brown University. This work was presented in part at the American Association for Cancer Research (AACR) annual meeting in Orlando, FL, April, 2023. The authors thank Robert Sobol, Sendurai Mani, and Bert Vogelstein for reading the manuscript and providing helpful suggestions. The authors thank Erika Tavares and Elizabeth L. Brainerd at the Keck XROMM Core Facility, Department of Ecology, Evolution and Organismal Biology, Brown University for assistance with CT imaging of mice.

## Author contributions

Conceptualization: W.S.E-D.; Methodology: W.S.E-D., J.L., A.L., M.H., P.S., A.G., A.D.L.C, L.Z. (Zhang), L.H.B., K.E.H., L.Z. (Zhou); Investigation: W.S.E-D., J.S., A.L., M.H., P.S., A.G., A.D.L.C, L.Z. (Zhang), L.H.B., K.E.H., L.Z. (Zhou); Visualization: W.S.E-D., J.L., A.L., M.H., P.S., A.G., A.D.L.C, L.Z. (Zhang), L.H.B., K.E.H., L.Z. (Zhou); Funding acquisition: W.S.E-D., A.E.A.; Project administration: W.S.E-D.; Supervision: W.S.E-D.; Writing – original draft: W.S.E-D.; Writing – review & editing: All authors.

## Data and materials availability

All data is included in the current manuscript. Genetic strains of mice that were used are commercially available.

---

**Table S1.** **Method used for 3D reconstruction of lung volume in CTAn software (Bruker) A)** Operations listed using terminology for CTAn that were utilized to reconstruct the lung volume of each individual mouse.

| Plug-in | Parameters | Result |
| --- | --- | --- |
| Thresholding | Global 14-255* | Segment body from air in lungs |
| Despeckle | 2D space, remove pores, by image borders, image | Remove the inside pores to generate a solid body |
| Despeckle | 2D space, sweep, all except the largest object, image | Remove the noise and the bed |
| Bitwise Operations | ROI = COPY image | Copy image to ROI |
| Reload | Image | Reload original image |
| Thresholding | Global 0-14* | Segment out the lungs |
| Bitwise Operations | Image = Image and ROI | Image now only contains white pixels belonging to both the image and ROI |
| Despeckle | 3D space, sweep, all except the largest object, image | Image only contains lungs in 3D space, removes fat and outside body |
| Despeckle | 3D space, remove black speckles, volume less than 33 voxels, image | Clean up the image and remove excess black pixels inside lung |
| 3D Analysis | Basic values, save results as text table | Excel sheet with volume of lungs |
| 3D model | *.ctm, Double-Time cubes | Builds and saves a 3D model file that can be opened in CTVol |

## Supplementary Figure Legends

**Figure S1.**
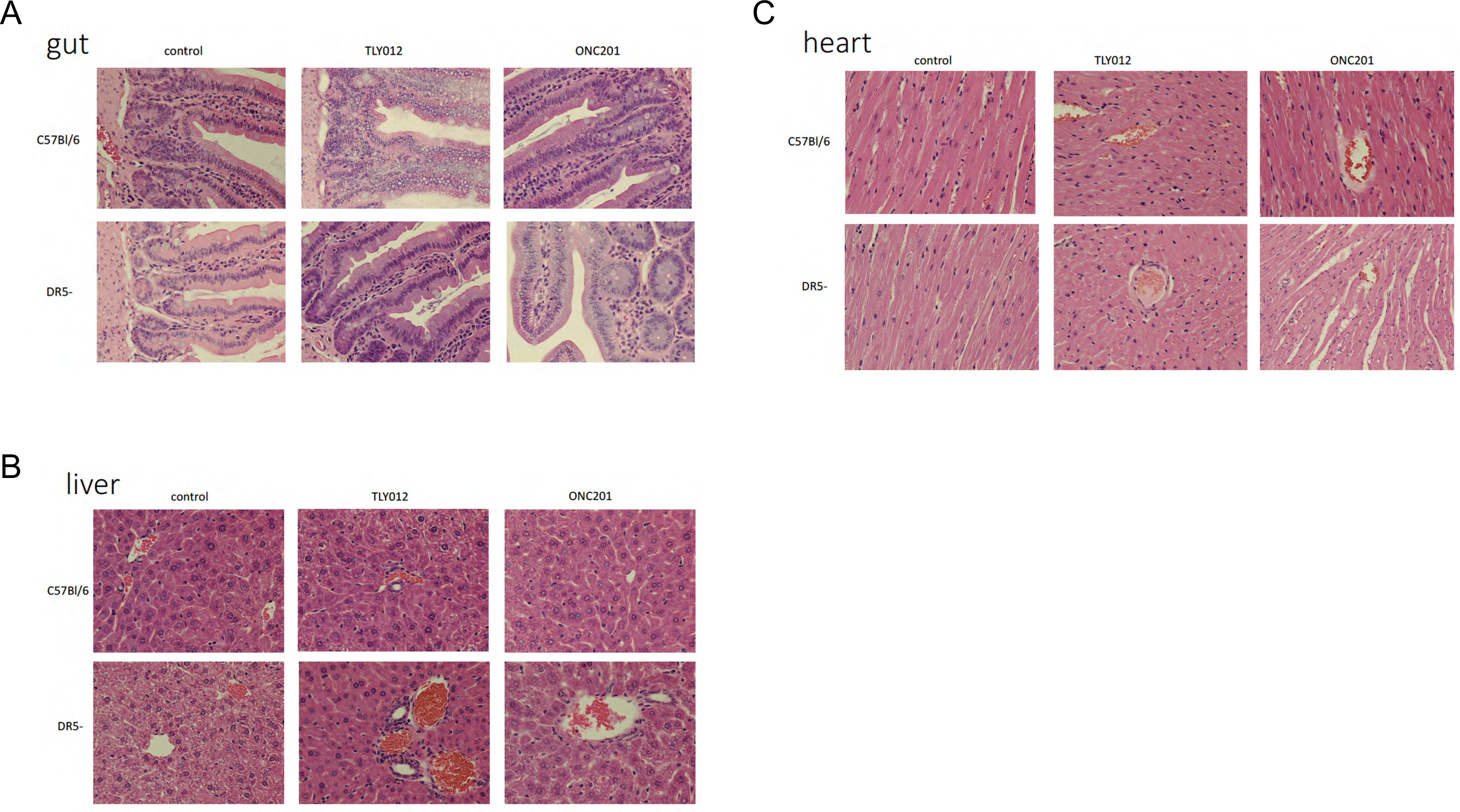
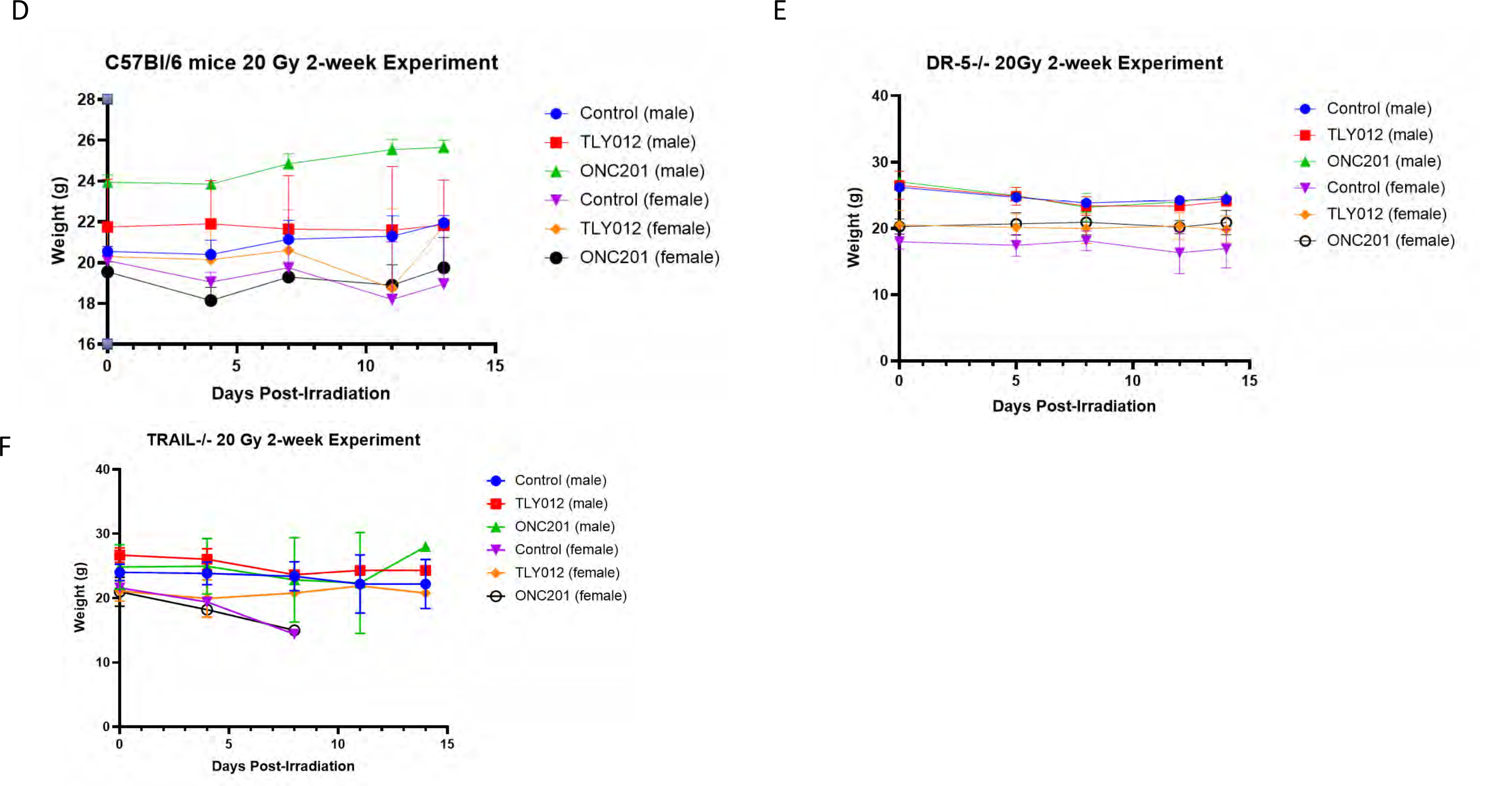
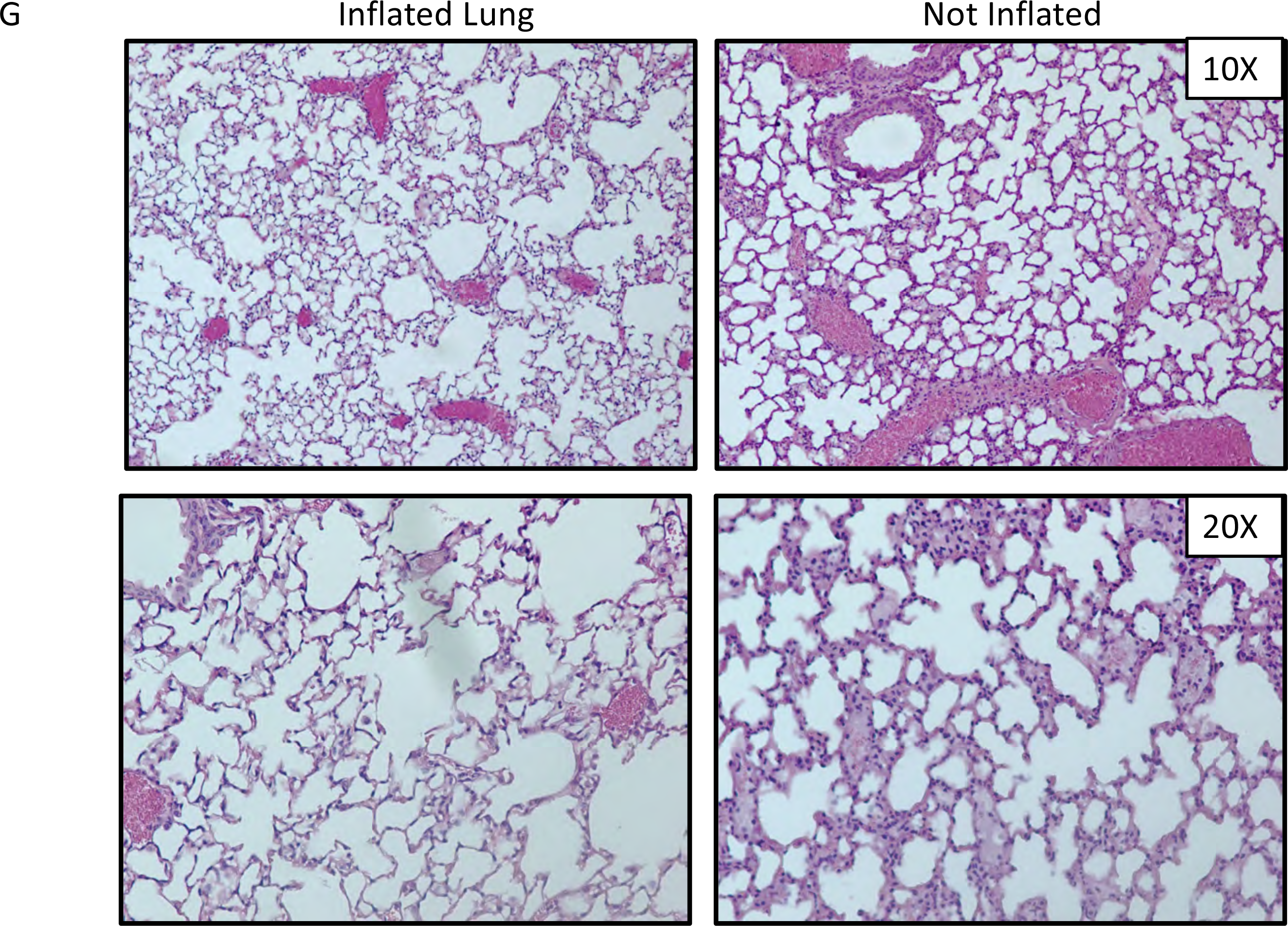
Radiation damage in other organs in the thorax. Lack of toxicity to duodenum (A), liver (B), and heart (C) after radiation (only chest radiation was administered). **D, E, F)** average of mouse whole body weights for the duration of the two weeks post whole-thorax x-ray irradiation of 20 Gy for C57Bl6, *DR5*-/-, and *TRAIL*-/- mice respectively (n = 2/gender/genotype/treatment). **G)** Lungs were not re-inflated post-mortem as determined by H&E-stained lung tissue at x10 and x20 inflated versus not inflated did not indicate major differences (n = 3 inflated, n = 1 not inflated).

**Figure S2.**
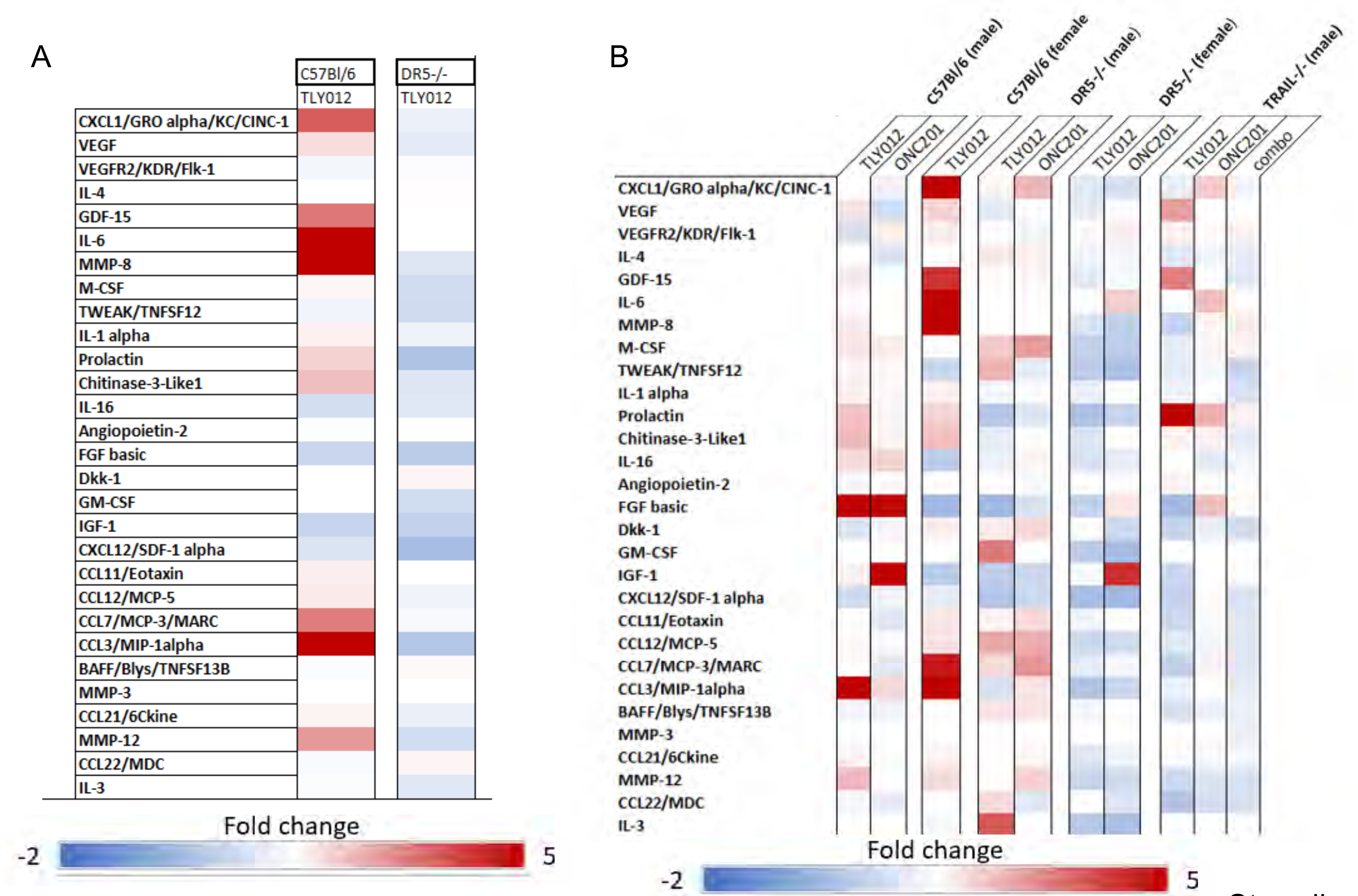
Cytokine alterations after radiation and TRAIL pathway agonist treatment in male and female mice. A) Heat map of TGF-β fold-change level 13 days post-irradiation compared to irradiated control (n = 2). **B)** Heat map of cytokine level fold change compared to irradiated control at 13 days post-irradiation (n = 2).

**Figure S3.**
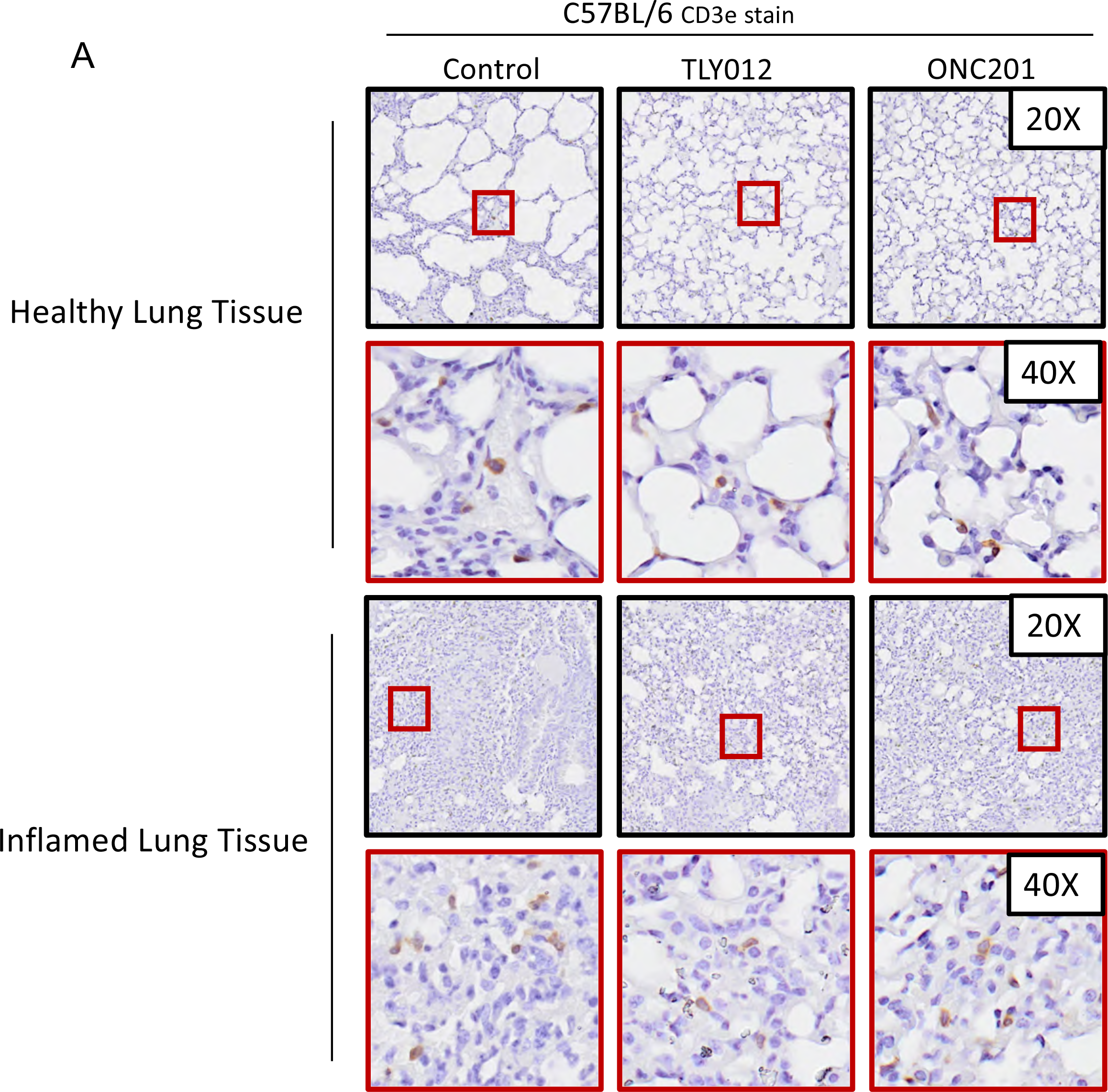
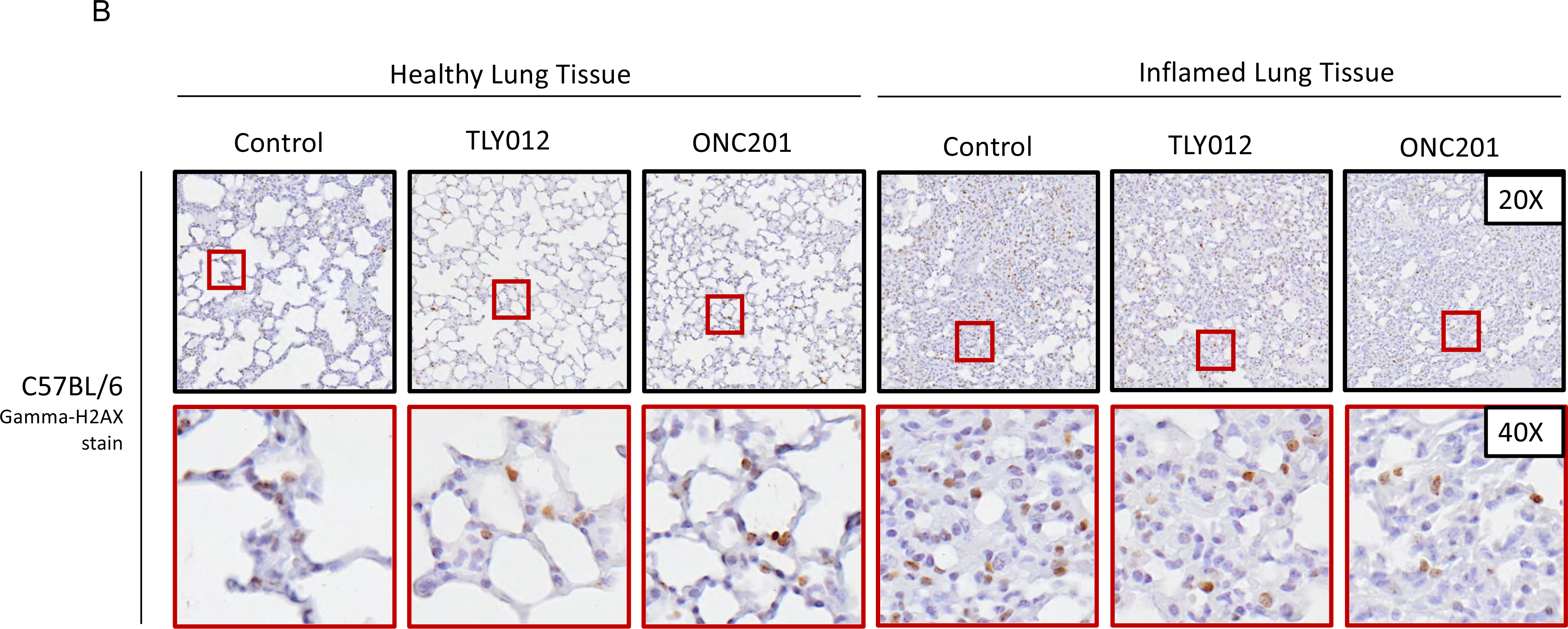
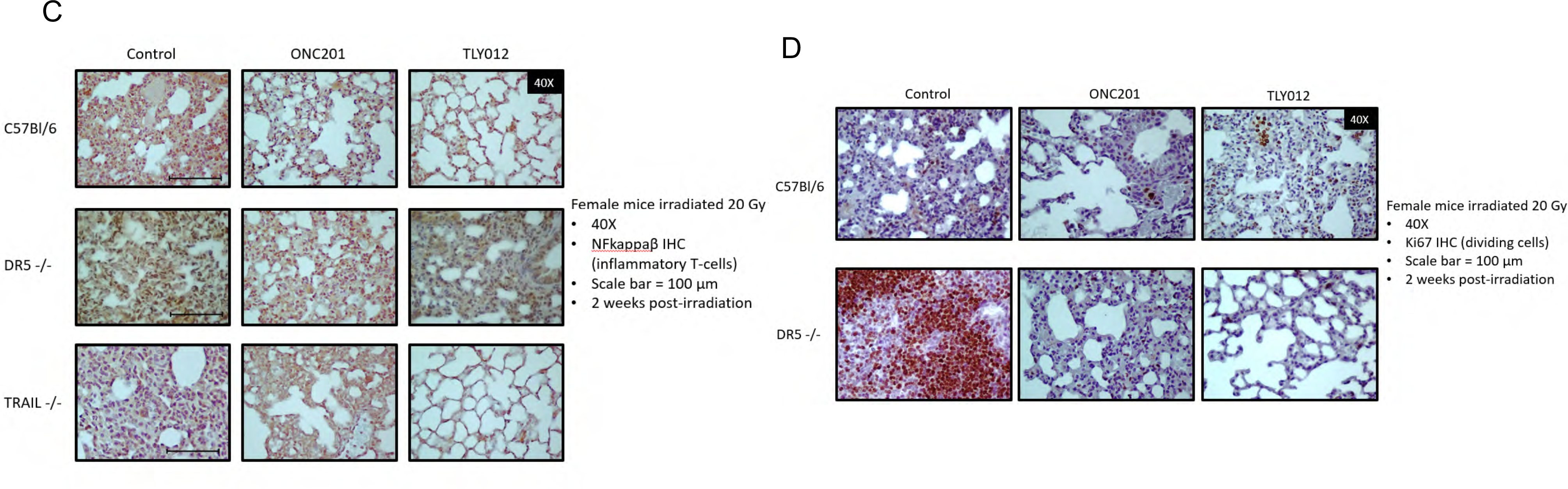
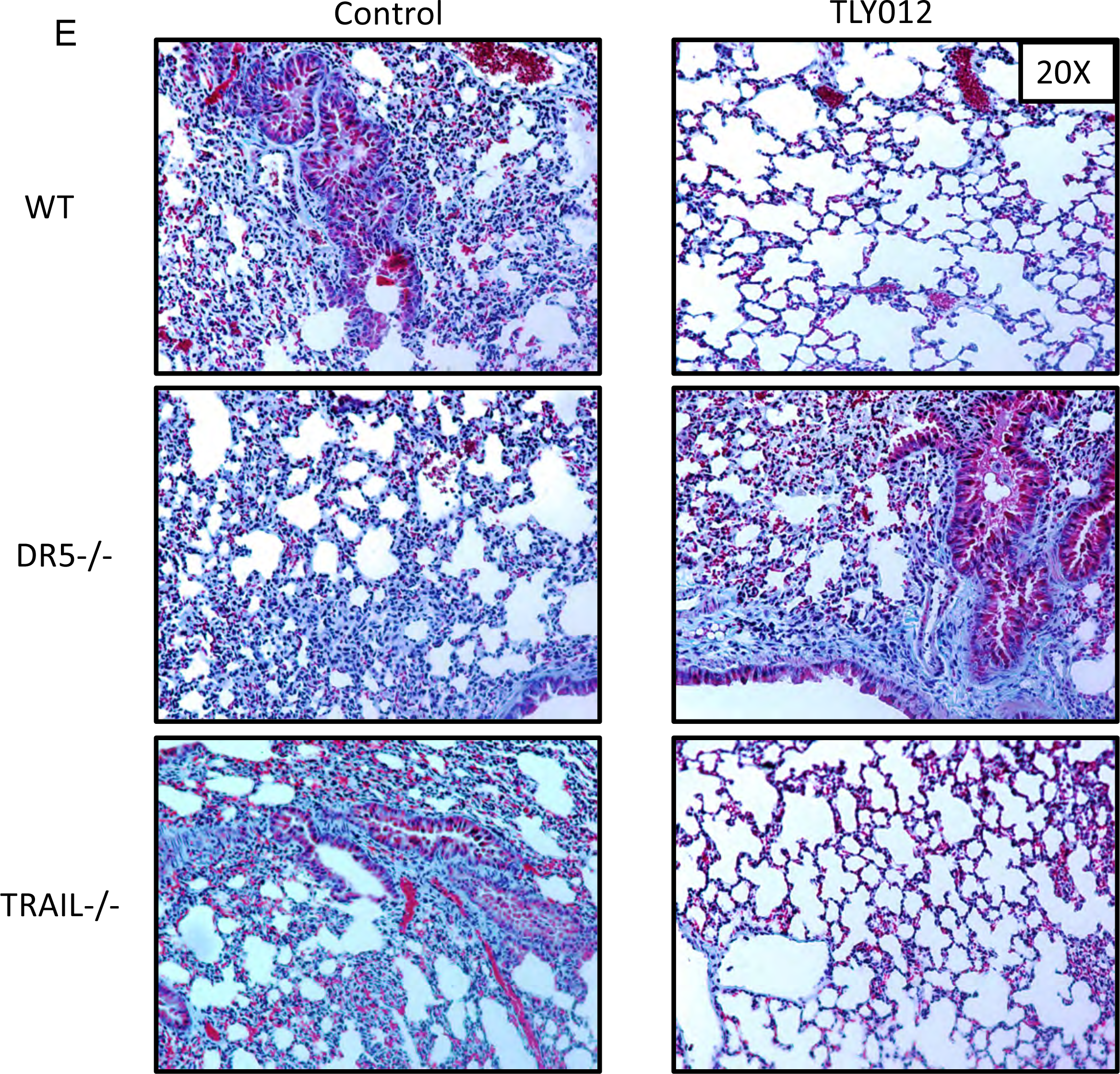
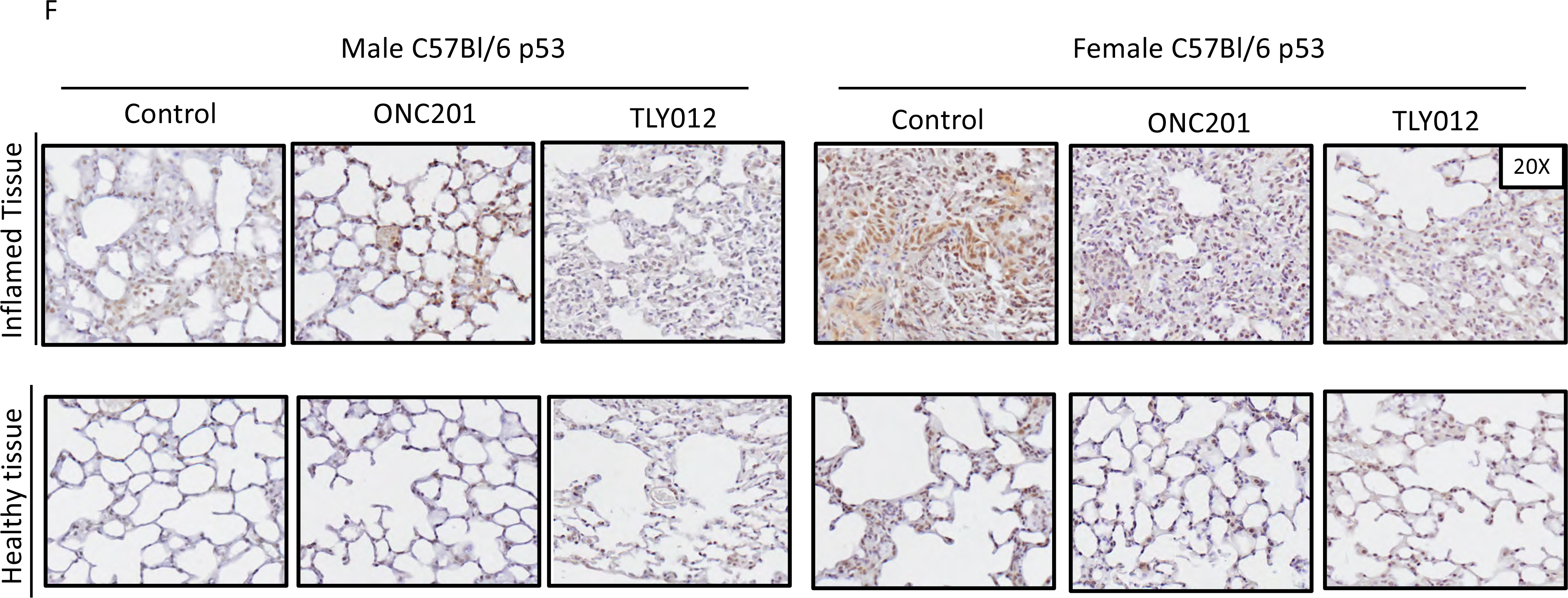
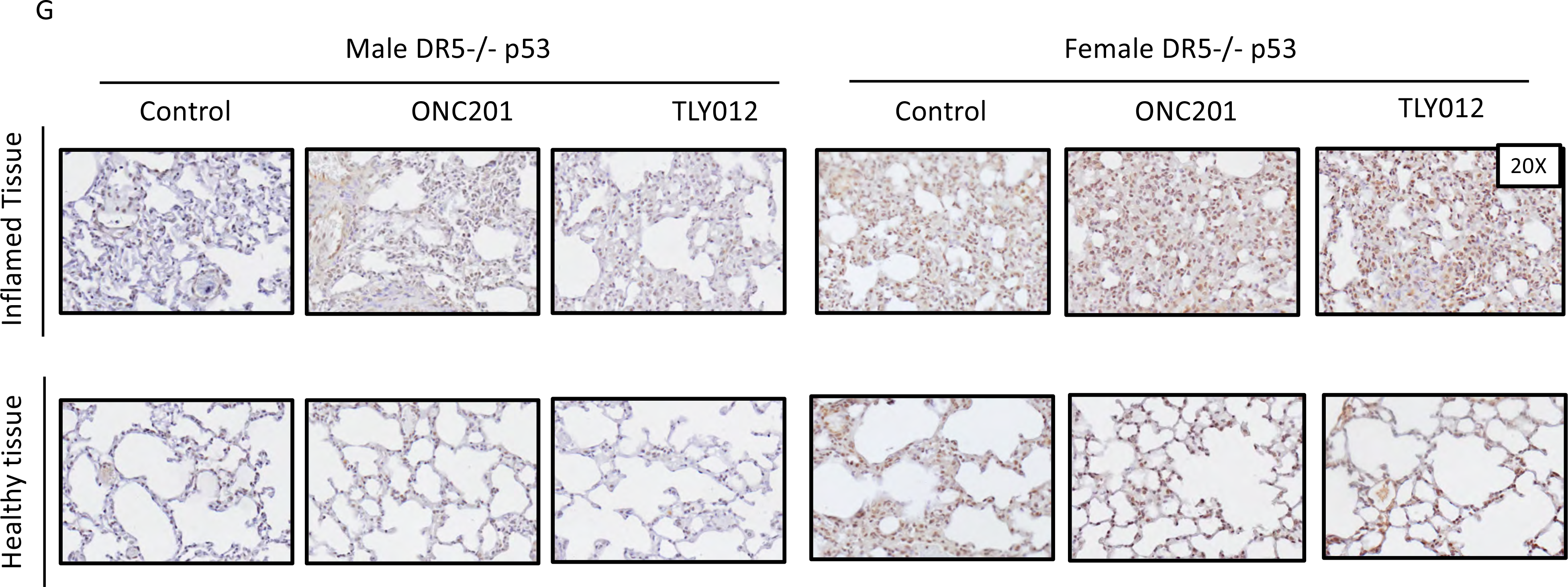
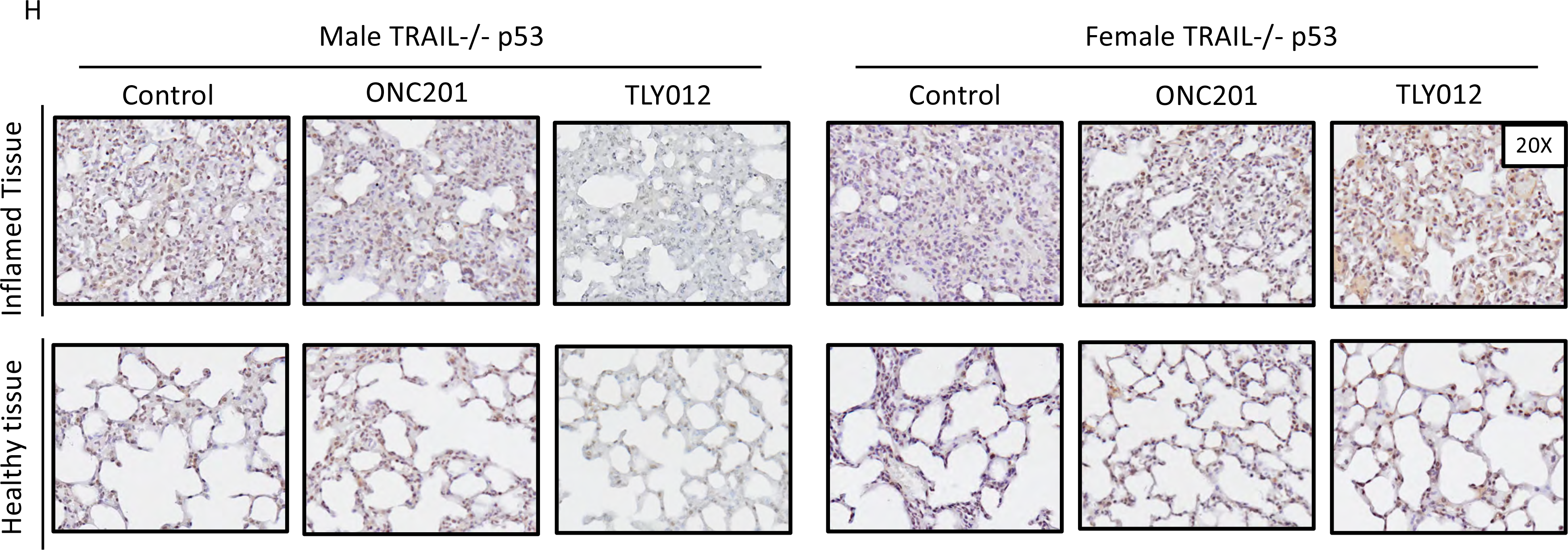
Immunohistochemical analysis of lung tissue irradiated with a single dose of 20 Gy x-ray irradiation in mice in C57Bl/6, *DR5*-/-, and *TRAIL*-/- genetic backgrounds +/- rescue with TRAIL pathway agonists. A) C57Bl/6 mice given a single whole-thorax irradiation dose of 20 Gy and treated with TLY012, ONC201, or control gavage were euthanized on day 13 post-irradiation. Immunohistochemical analysis of CD3ε was performed to determine T-cell infiltration. **B)** Mice treated with TLY012 showed a decrease in gamma-H2AX in both inflamed and healthy areas of lung tissue (top row x20, bottom row x40). **C)** Immunohistochemistry of female mice shows abundant infiltration of NFkB in all three genotypes C57Bl/6, *DR5*-/-, and *TRAIL*-/- in all treatment groups. Scale bar: 100 μm. **D)** Immunohistochemistry shows abundant labeling of the proliferation marker Ki67 (DAB, brown staining) in *DR5*-/- control mice at 13 days-post whole-thorax irradiation of 20 Gy. **E)** Trichrome staining showed an increase in collagen deposition (light blue) in WT and *TRAIL*-/- control mice compared to those rescued with TRAIL agonist. **F, G, H)** p53 in male and female mice with C57Bl/6, *DR5*-/-, and *TRAIL*-/- backgrounds respectively. Areas of inflamed tissue and healthy tissue were examined at 20X.

**Figure S5.**
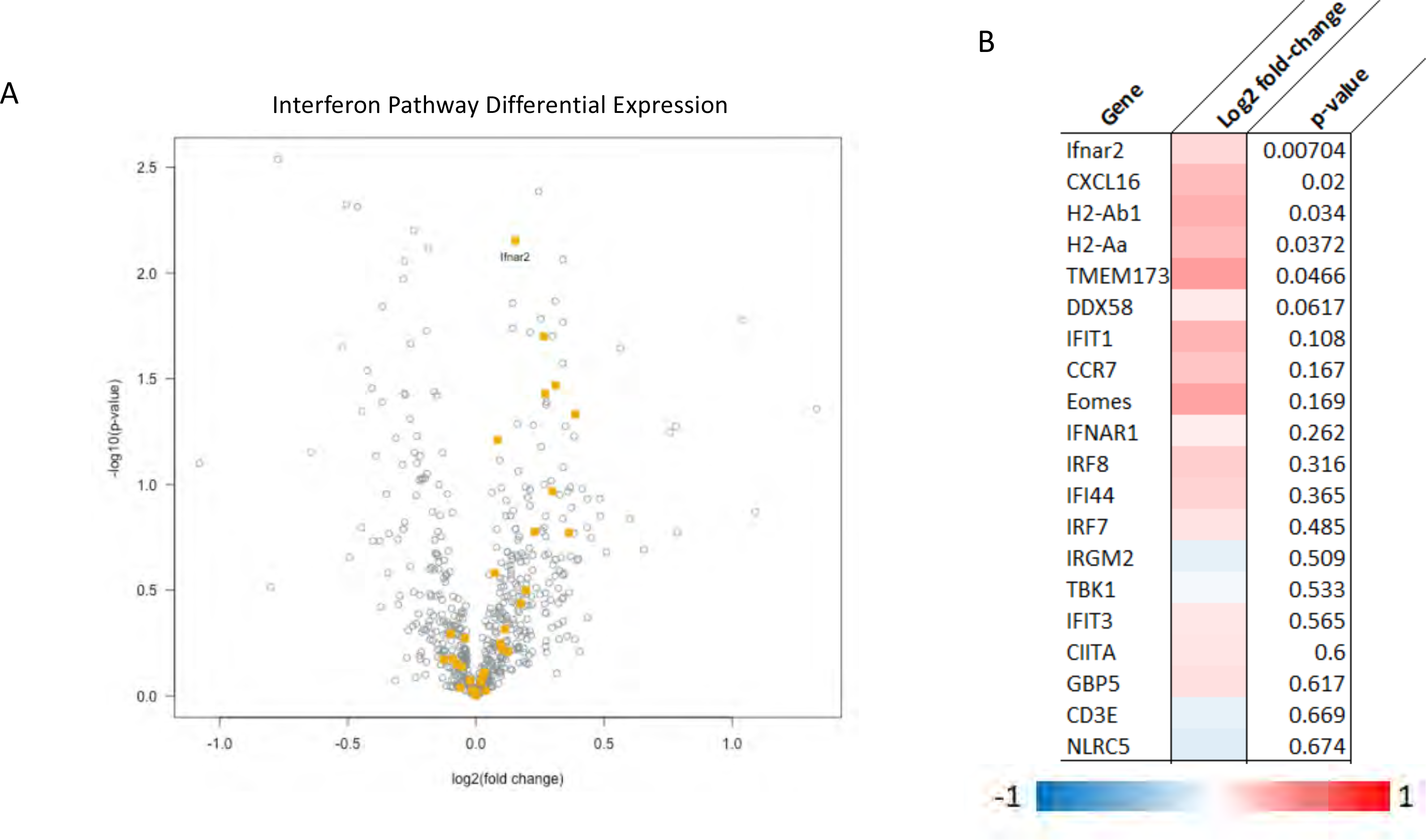
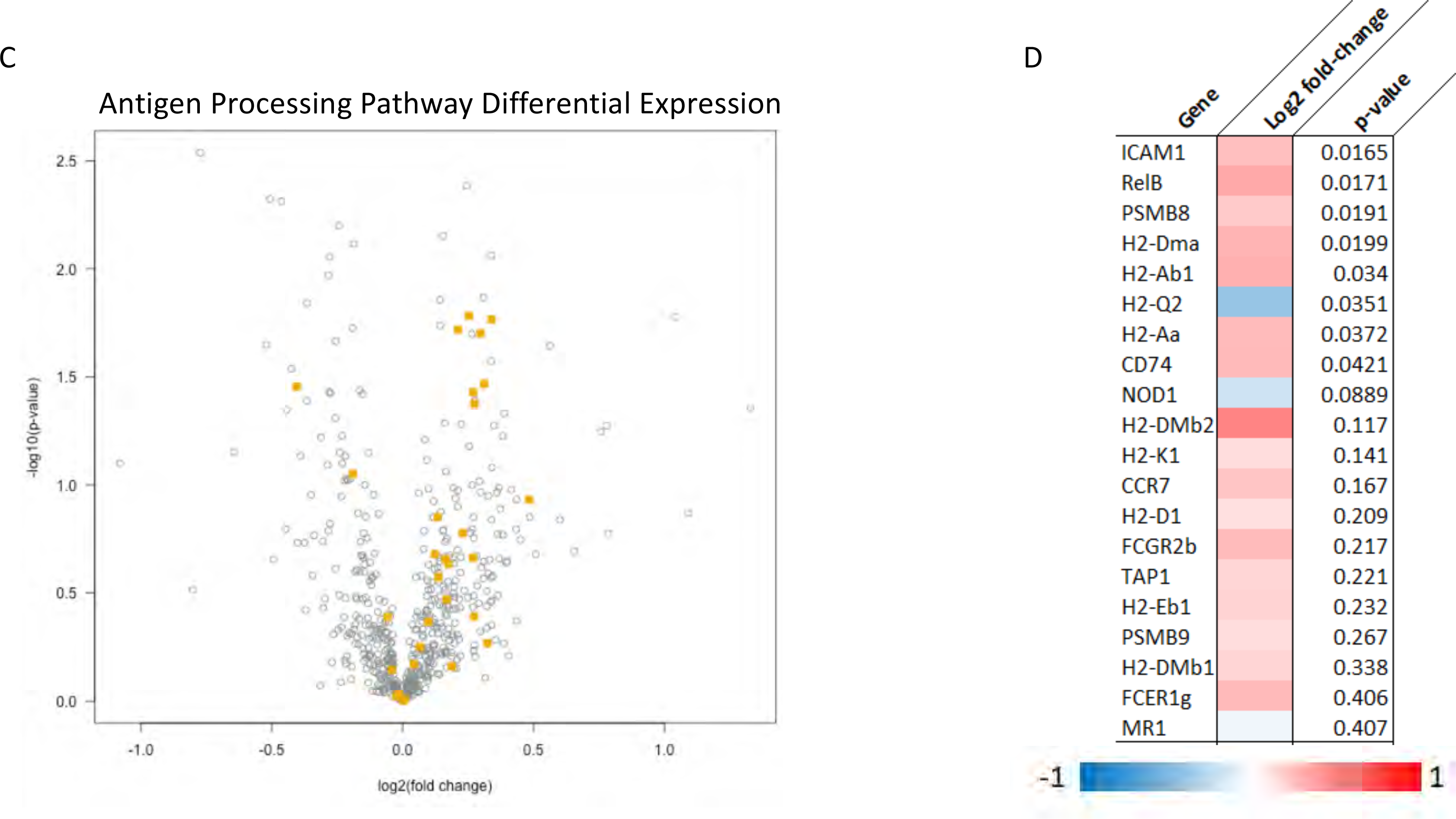
Immune pathways were significantly impacted by TLY012 treatment. A) Volcano plot of interferon pathway genes with significant differential expression in the TLY012 treated group compared to control (n = 6/treatment/group). **B)** Heat-map of the fold-change of genes in the interferon pathway and their statistical significance. **C)** Volcano plot of antigen processing pathway genes with significant differential expression in the TLY012 treated group compared to the control (n = 6/treatment/group). **D)** Heat-map of the fold-change of genes in the antigen processing pathway and their statistical significance.

**Figure S6.**
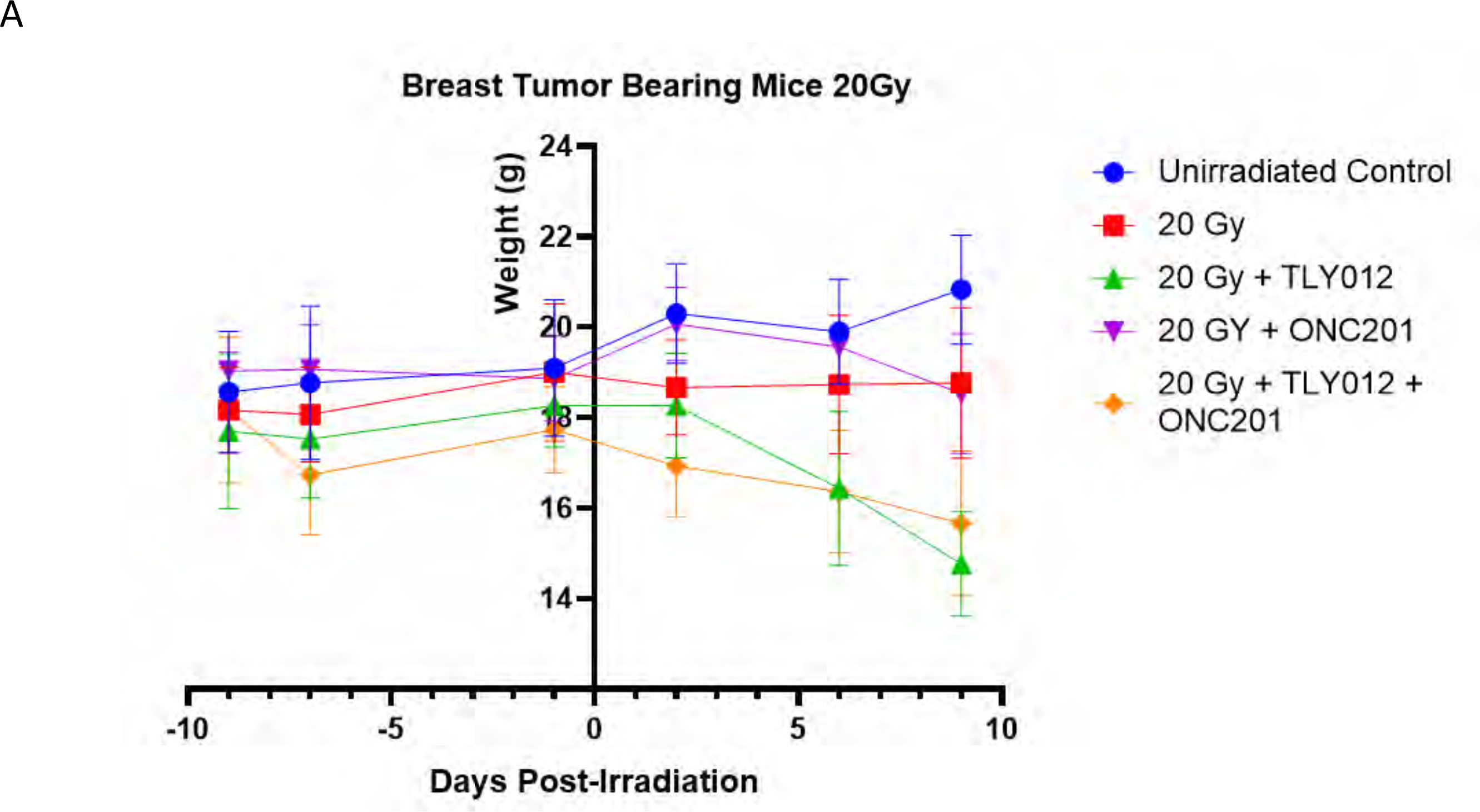
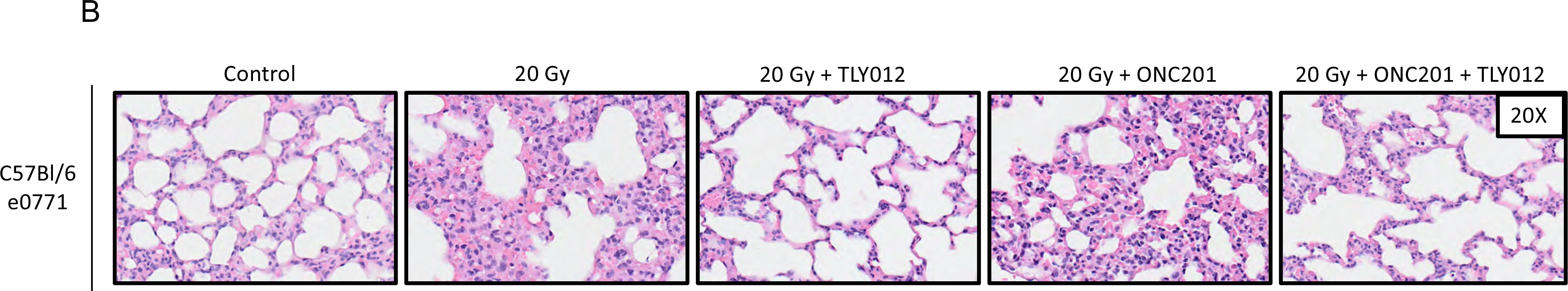
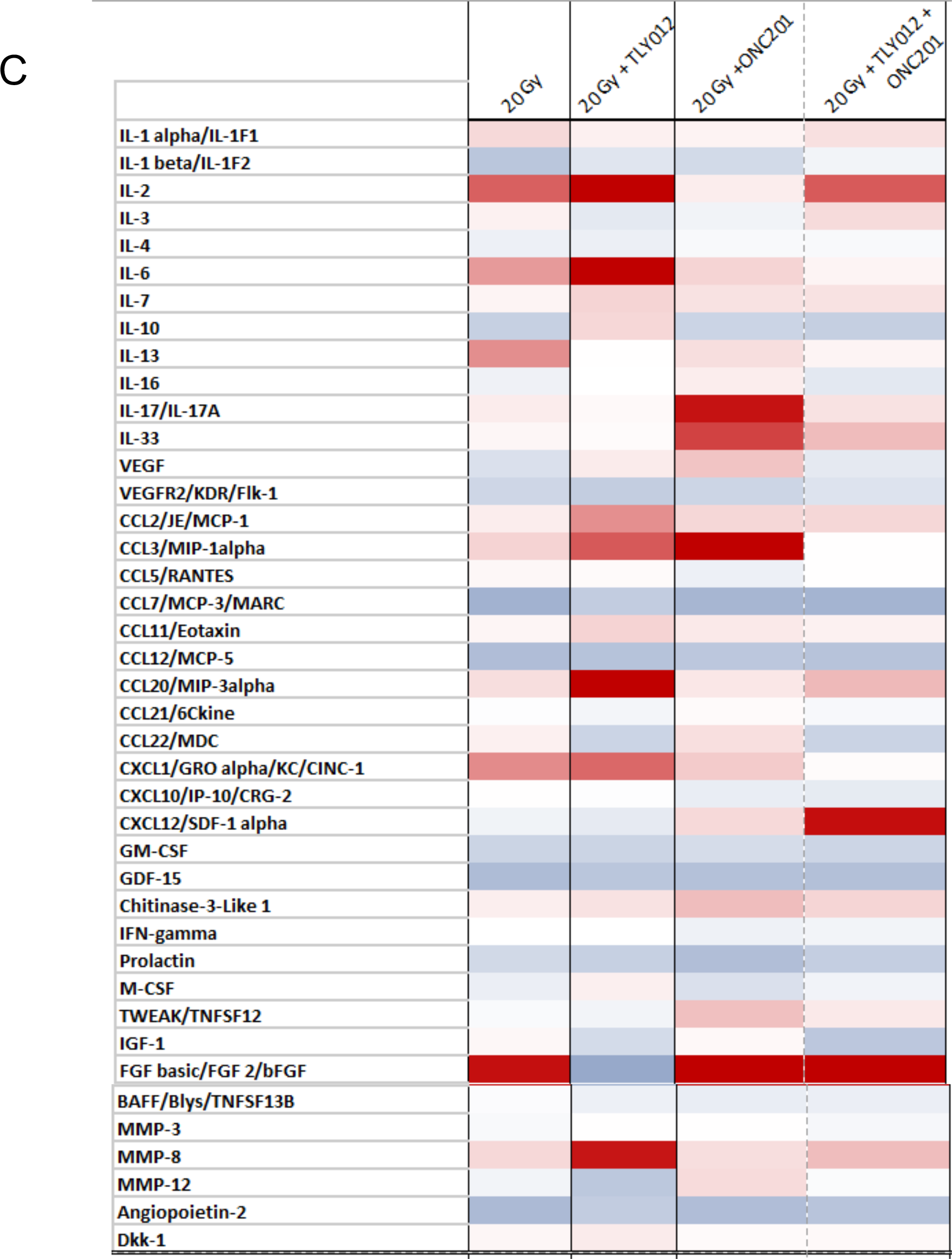
Histological and cytokine alterations in female immune competent C57Bl/6 mice with breast tumor xenograft -/+ rescue with TRAIL pathway agonists. A) Average total body weight of mice from orthotopic tumor injection (day = -9), through radiation (day = 0), up until sacrifice on day 9 post-irradiation (n=3/treatment/group). **B)** H&E stains of lung tissue at 9 days-post irradiation showed a reduction in alveolar-wall thickness and decreased inflammation in mice treated with TLY012. **C)** Heat-map of cytokine level fold-change of treated mice to unirradiated control.

**Figure S7.**
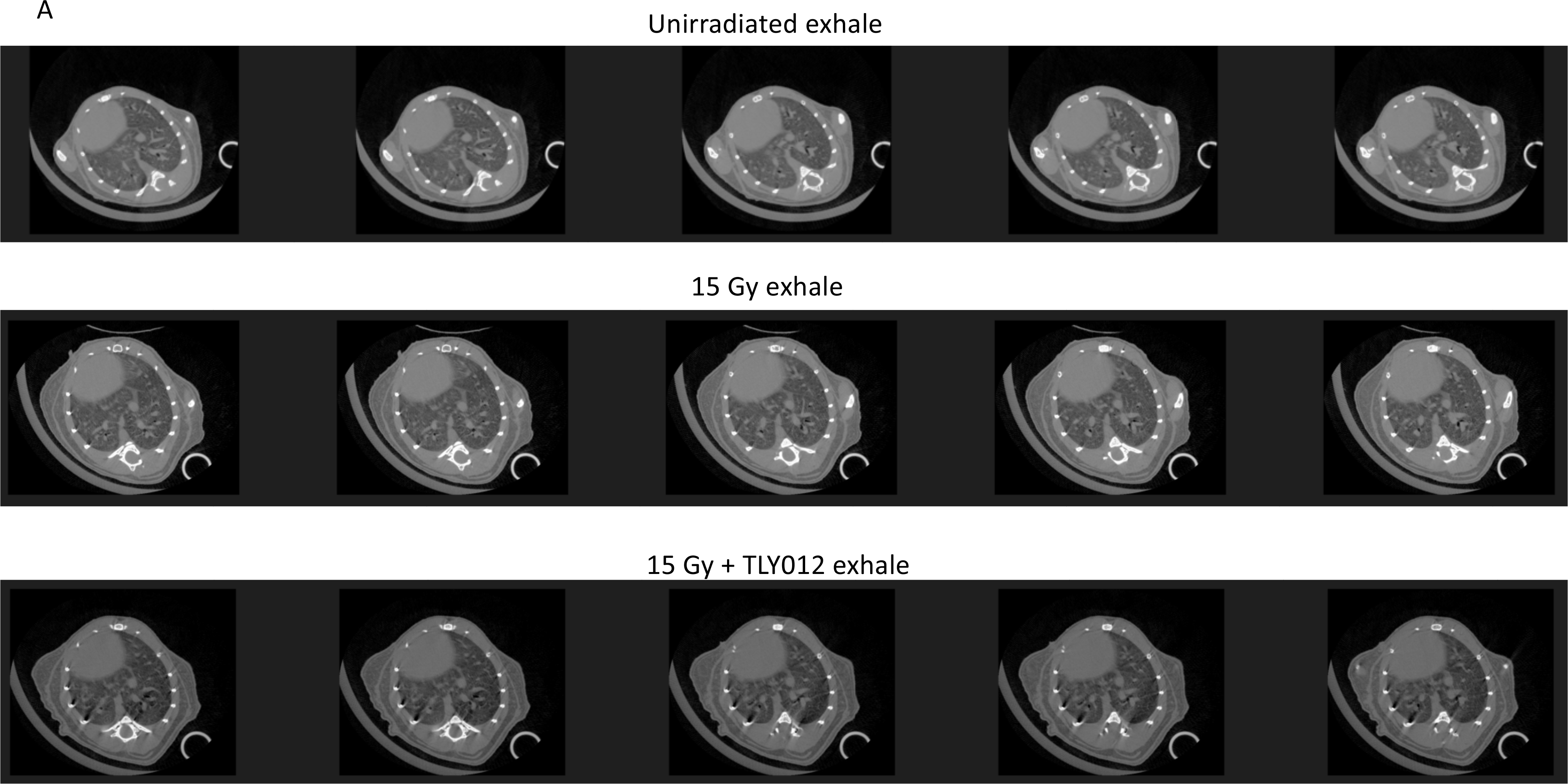
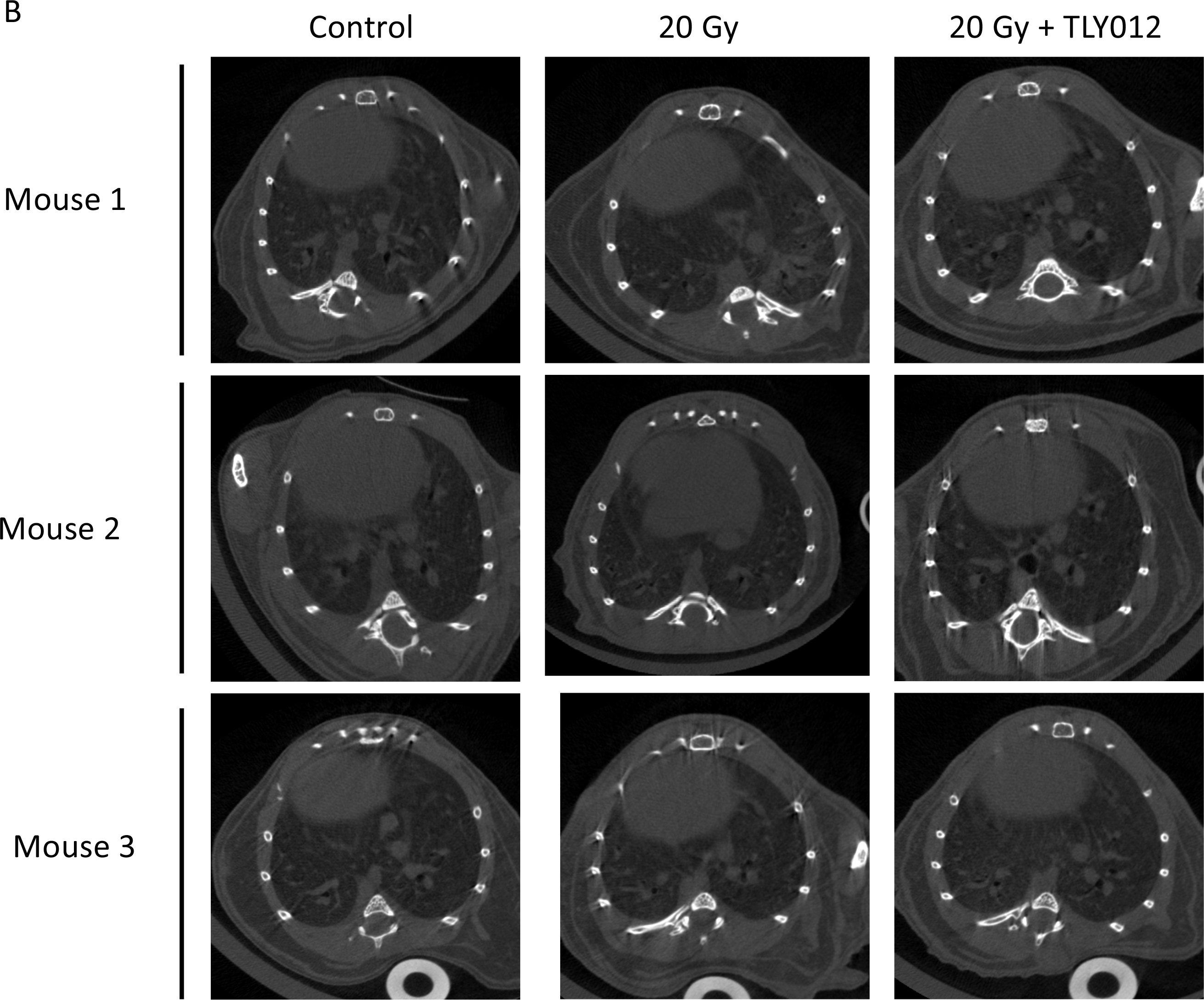
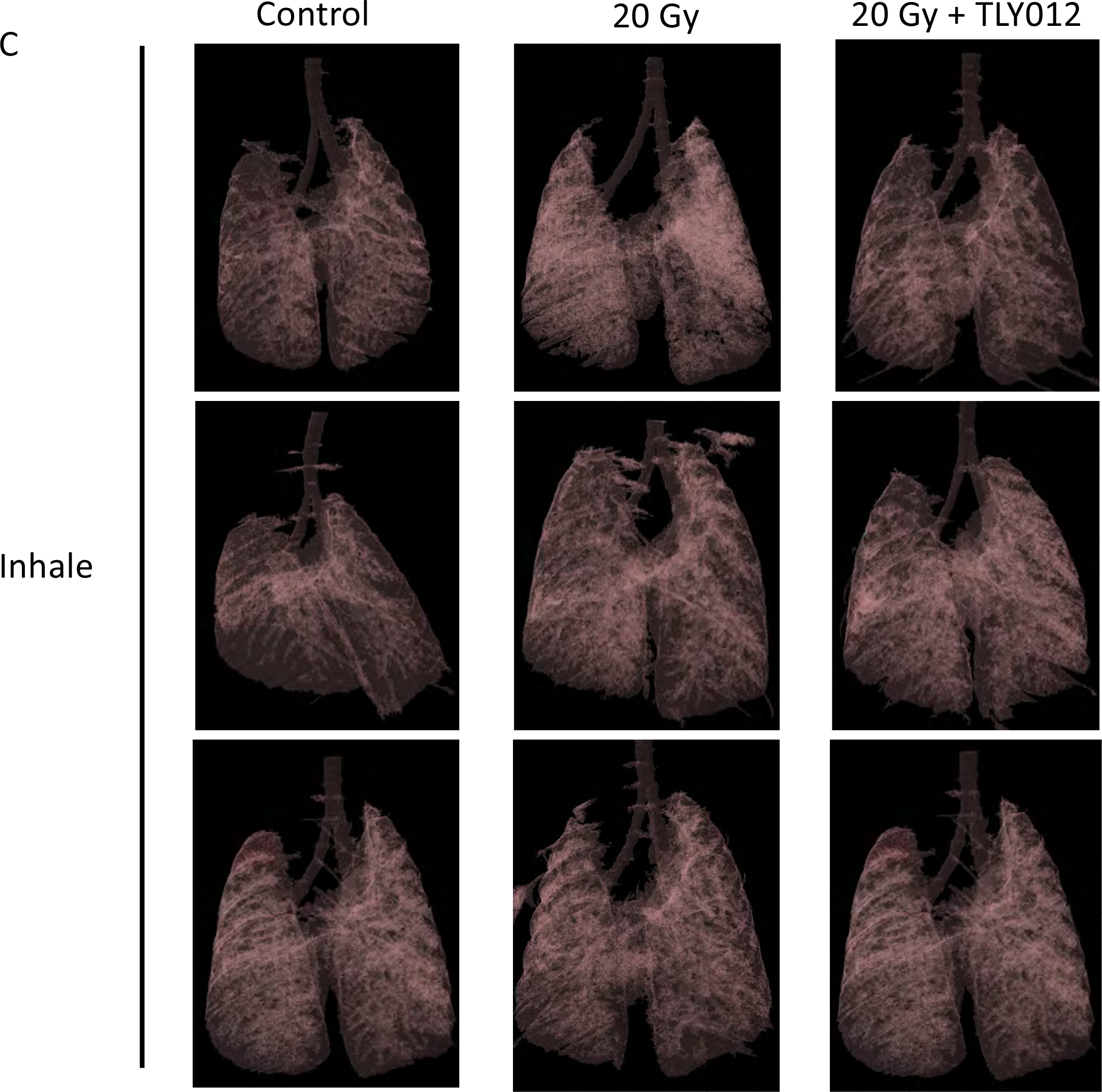
Additional representative μCT images of mice unirradiated and irradiated with 15 Gy whole-thorax x-ray irradiation and treated with or without TLY012. A) μCT images of the exhale duration of breathing cycle of mice that were unirradiated (top row), received one whole-thorax x-ray irradiation dose of 15 Gy (middle row), and received 15 Gy irradiation in addition to rescue with twice a week treatment of TLY012 for two weeks (bottom row). **B)** μCT images of the inhale duration of the breathing cycle in mice that were unirradiated, received one whole-thorax x-ray irradiation dose of 20 Gy, and received 20 Gy irradiation in addition to rescue with twice a week treatment of TLY012 (n = 3/treatment/group). **C)** 3D reconstruction of the μCT images during exhale and inhale portions of the breathing cycle. All images are subjected to a 7% opacity filter in CTVol software.

**Figure S8.**
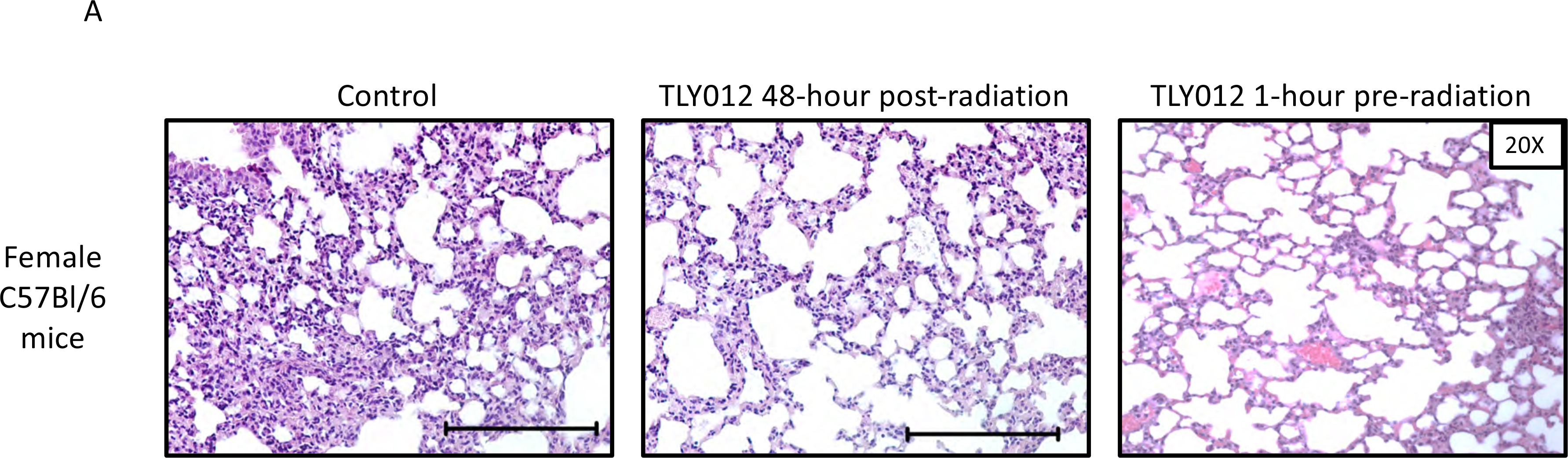
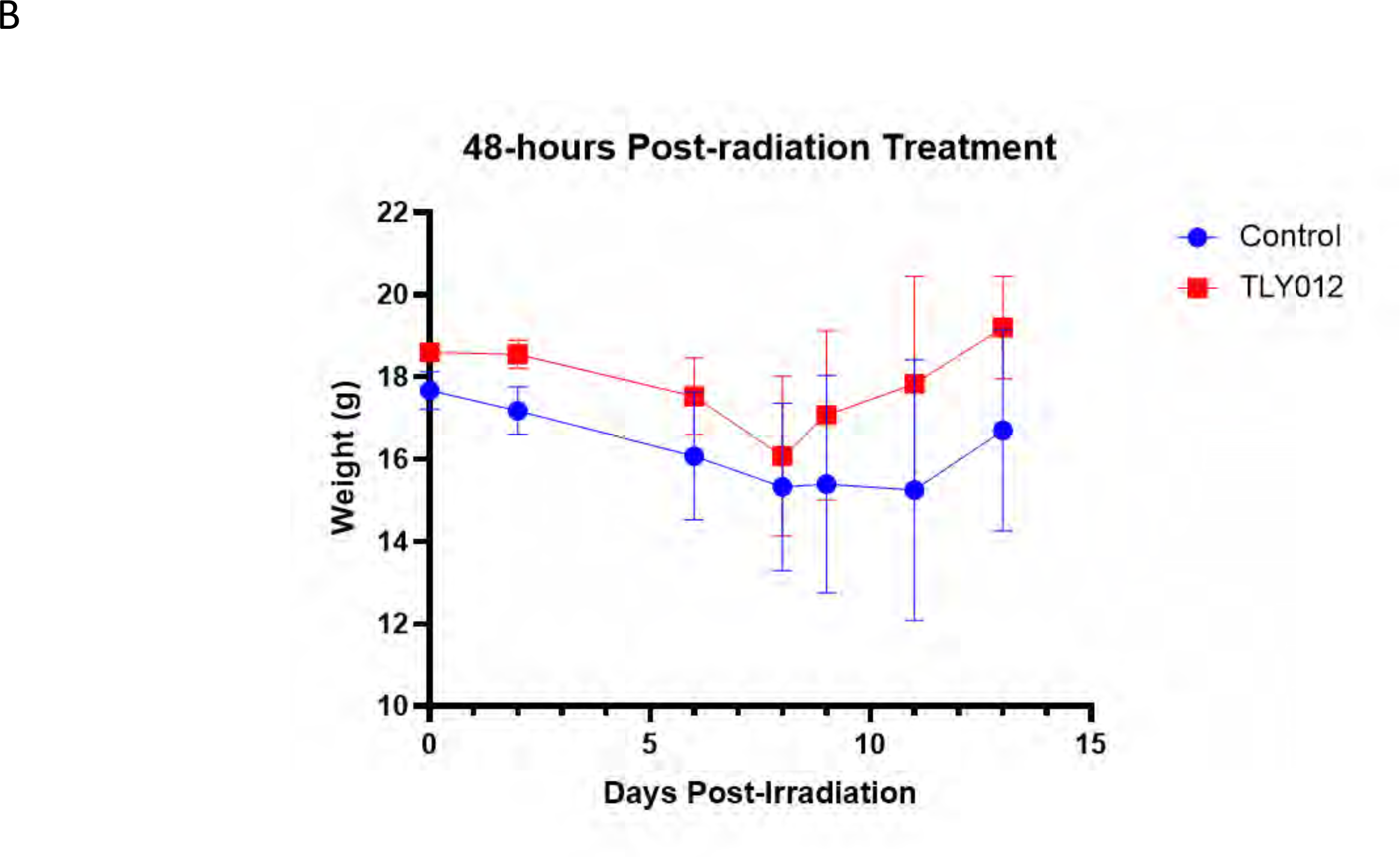
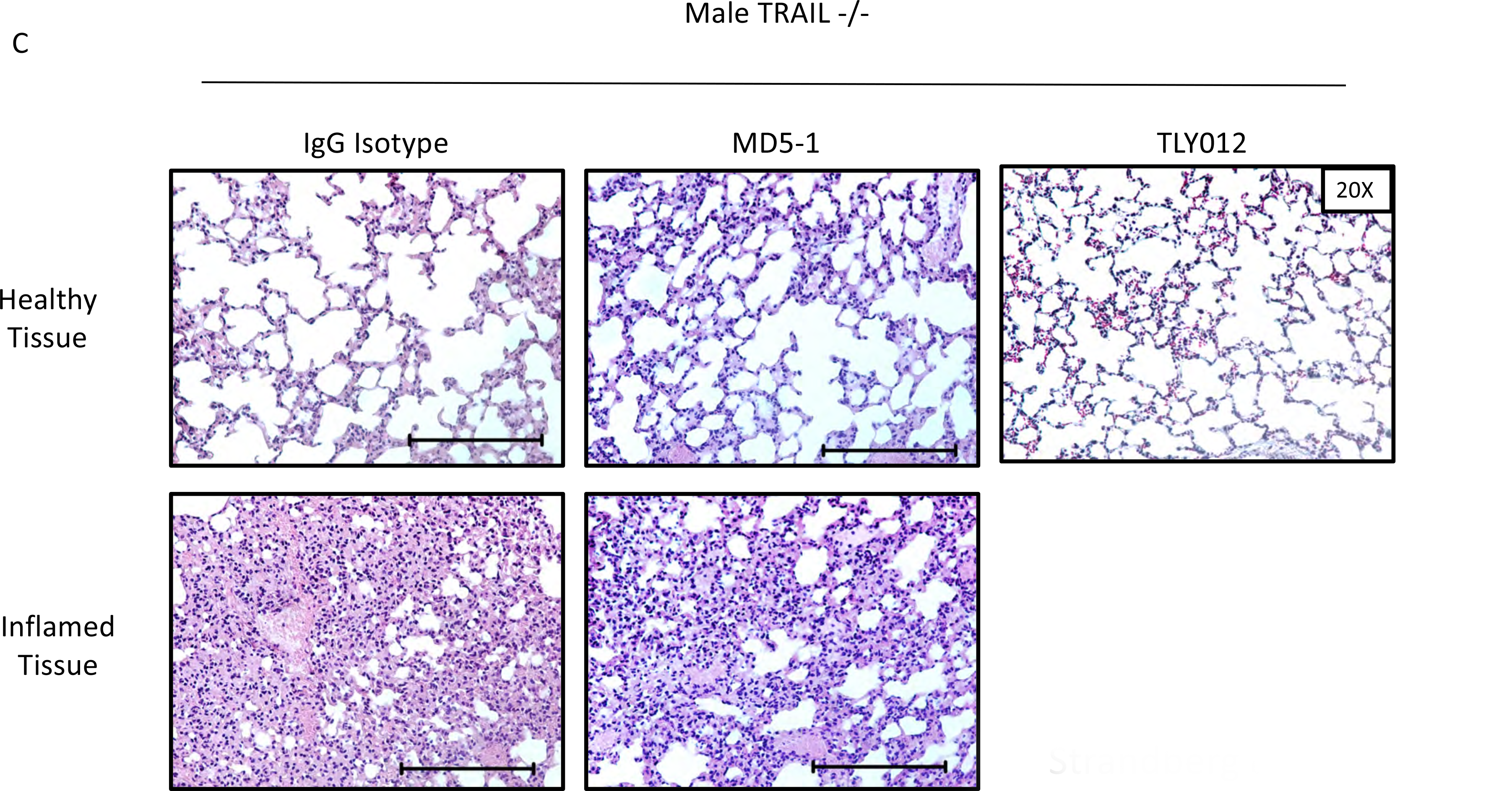
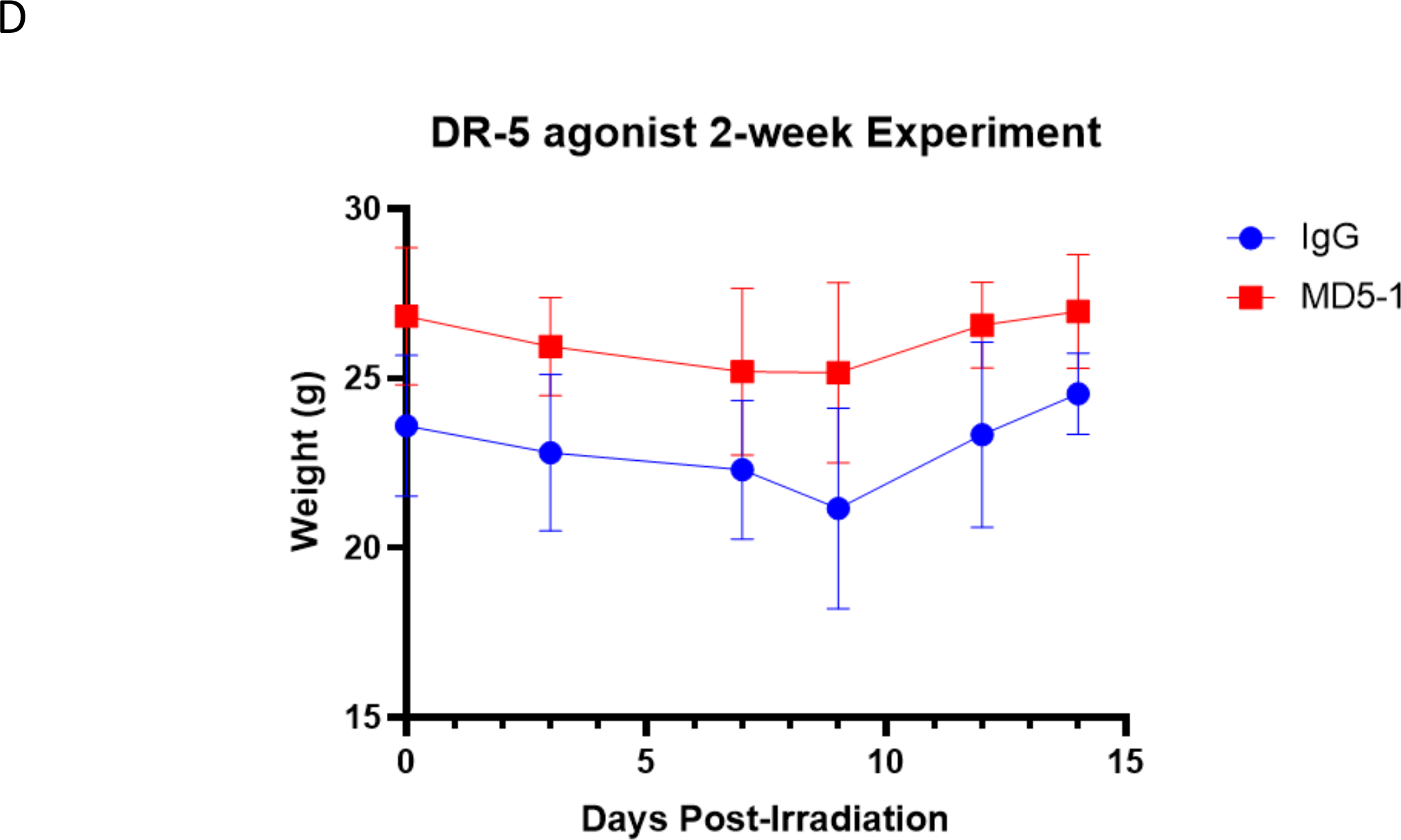
Additional TRAIL agonist experiments including 48-hour post-irradiation treatment with TLY012. A) 48-hour post whole-thorax x-ray irradiation TLY012 was administered continuing twice a week. **B)** Average full body weights of control mice or mice that began twice a week treatment of TLY012 48 hours after receiving one whole-thorax x-ray irradiation dose of 20 Gy (n = 3/treatment/group) **C)** Male *TRAIL*-/- mice treated with antibody MD5-1 or control IgG isotype once a week. **D)** Mice treated with MD5-1 or IgG isotype average whole-body weights for two weeks of treatment post-radiation (n = 3/treatment/group).

